# Evaluating Microglial Contributions to the Neurovascular Unit in Health and Neurodegeneration Using Human *In Vitro* Models

**DOI:** 10.64898/2025.12.19.695574

**Authors:** Kira Evitts, Emily Turschak, Charles A. Williams, Chizuru Kinoshita, Aleah Rosner, Willow Battista, Katherine Hui, Ian Beck, Aquene Reid, Ying Zheng, Jessica E. Young

**Affiliations:** Department of Bioengineering; University of Washington; Seattle, WA, 98109; United States of America; Institute for Stem Cell and Regenerative Medicine, University of Washington, Seattle, WA 98109, United States of America; Department of Laboratory Medicine and Pathology, University of Washington, Seattle, WA 98109, United States of America

**Keywords:** Neurovascular unit, Alzheimer’s Disease, iPSC-derived microglia, 3D engineered microvessels, inflammation, brain endothelial cells

## Abstract

**Background:** Microglia are emerging as critical regulators of neurovascular function in health and Alzheimer’s disease (AD), yet their interactions with the human neurovascular unit (NVU), particularly brain endothelial cells, remain incompletely understood. Current *in vitro* NVU platforms typically exclude microglia and lack perfusable vascular networks with physiologically relevant architecture. Here, we established complementary two-dimensional (2D) and three-dimensional (3D) NVU models to investigate microglia-endothelial and microglia-neurovascular interactions.

**Methods:** Human induced pluripotent stem cell derived-neurons (iNs), astrocytes (iAs), and microglia-like cells (iMGLs) were incorporated into a soft-lithography based engineered microvessel system to establish a multicellular neuroimmune-vascular model. To specifically evaluate iMGL-endothelial cell (EC) interactions, iMGL were co-cultured with primary human brain microvascular endothelial cells (HBMECs) and junctional protein localization was evaluated using immunofluorescence. The barrier integrity of engineered microvessels containing iMGL was evaluated using dextran permeability. Our 2D and 3D systems were stimulated with tumor necrosis factor-α (TNFα) to evaluate whether iMGL would promote or attenuate EC inflammation and barrier breakdown.

**Results:** Incorporation of iNs, iAs, and iMGLs into a perfusable vascular model enabled a more complete representation of NVU cellular diversity and promoted neuronal health. In monolayer co-culture with iMGL, HBMECs enhanced the junctional localization of tight and adherens junction proteins through both contact-dependent and paracrine mechanisms. Following an inflammatory challenge, iMGLs reduced endothelial inflammatory activation, suggesting a protective role in response to AD-relevant inflammatory conditions. Finally, when embedded in 3D collagen matrices surrounding perfusable endothelialized lumen networks, iMGLs reduced dextran permeability and preserved endothelial barrier integrity following TNFα challenge.

**Conclusions:** Together, these findings establish a 3D perfusable neuroimmune-vascular model that enables the dissection of microglial contributions to neurovascular function in health and disease.

## 1. Background

The human neurovascular unit (NVU) is a specialized multicellular system that ensures adequate cerebral blood flow and nutrient delivery to support neuronal health^1, 2^. The NVU is composed of neurons, astrocytes, microglia, pericytes, and specialized brain microvascular endothelial cells that form the blood-brain barrier (BBB)^3–5^. The BBB maintains central nervous system (CNS) homeostasis by tightly regulating paracellular and transcellular transport of ions, molecules, and cells. This selective barrier function is mediated by tight-junction proteins such as claudin-5, occludin, and zonula occludens, which link endothelial junctions to the actin cytoskeleton^4–6^.

Vascular dysfunction is known to contribute to Alzheimer’s disease (AD)^7–11^. AD is associated with compromised NVU and BBB function, which promotes vascular leakage, neuroinflammation, and accumulation of neurotoxic molecules, thereby accelerating neuronal loss and cognitive impairment ^1, 8, 9, 11–14^. The dysfunction of BBB is thought to emerge early, typically before or during mild cognitive impairment and initial symptomatic stages^12, 13, 15–21^. BBB breakdown in AD involves loss of tight-junction proteins and subsequent leakage of neurotoxic blood-derived proteins into the brain parenchyma^12, 13, 18–21^. This can lead to activation of microglia and astrocytes resulting in the release of pro-inflammatory cytokines, including interleukin (IL)-1β, TNFα, and IL-6, which in turn contribute to the neuronal degeneration observed in post-mortem AD brains^2, 19, 22–26^. Although NVU impairment in AD is well recognized, the precise cellular mechanisms by which individual NVU cell types contribute to dysfunction, and how NVU cell types interact during health and disease, remain incompletely understood.

Microglia are brain-resident immune cells with essential roles in maintaining CNS homeostasis and responding to pathology. In the healthy adult brain, they actively survey the microenvironment, refine neural circuits via synaptic pruning, clear apoptotic cells and protein aggregates through phagocytosis, and secrete neurotrophic and anti-inflammatory factors to support tissue integrity^27–30^. Genome-wide association studies (GWAS) have identified several microglial genes, including *TREM2*, *CD33*, *ABCA7, BIN1*, *SPI1*, and *PICALM,* as key genetic risk factors for late-onset AD, underscoring microglia’s central role in AD pathogenesis^31–38^. However, their dynamic, context-dependent activation states remain difficult to characterize^31^. While murine models have provided critical insights, species-specific differences in microglial gene expression, especially in aging and disease, highlight the need for human-based systems to better understand microglial behavior^39, 40^.

A range of *in vitro* human cell-based models have been developed to study the BBB and NVU, particularly in the context of AD. Traditional transwell-based co-culture systems incorporating endothelial cells, astrocytes, and pericytes allow for basic assessment of barrier integrity, such as transendothelial electrical resistance (TEER) and paracellular permeability^41–43^. While widely used, these static models lack physiological flow and structural complexity. More advanced microfluidic organ-on-a-chip platforms have improved upon these limitations by introducing dynamic shear stress and, in many cases, human induced pluripotent stem cell (iPSC)-derived NVU components to better model disease-relevant biology^44–51^. However, despite increasing complexity, most current models either omit microglia entirely or fail to capture their functional interactions with the brain vasculature.

The nature and extent of microglia-EC interactions in the NVU remain incompletely characterized. In the adult brain, it is estimated that approximately 20–30% of microglia associate with brain capillaries and form somatic contacts with ECs^52–54^. Most studies investigating EC-microglia interactions have focused on conditions of vascular injury or inflammation to better understand how microglia influence brain vasculature in inflammatory and disease contexts. For example, microglia have been shown to have a dual role in a mouse model of systemic inflammation where microglia initially preserve EC barrier integrity, but later contribute to its disruption as inflammation progresses^54^. In contrast, 2D *in vitro* co-culture studies have shown that microglia can impair endothelial barrier function in response to acute lipopolysaccharide (LPS) stimulation^55, 56^. These discrepancies highlight the elusive role of microglia in regulating EC barrier integrity.

In the healthy NVU, evidence regarding whether microglia support BBB integrity and tight junction expression remains limited and conflicting. An *in vivo* study using pharmacological microglial depletion for one month in mice reported no significant changes in BBB ultrastructure, gene expression, or permeability^57^. A separate study using Evans blue dye similarly found that elimination of microglia did not disrupt BBB integrity^58^. In contrast, *in vitro* models have demonstrated that co-culturing brain ECs with microglia or microglia-conditioned media increases protein levels of tight junction markers occludin and zonula occludens-1 (ZO-1)^59, 60^. Moreover, disruption of EC-microglia crosstalk via attenuated Colony-Stimulating Factor 1 Receptor (CSF-1R) signaling resulted in reduced claudin-5 expression *in vitro*^59^. Taken together, these findings suggest that microglia may influence the BBB under baseline conditions, but their role is difficult to discern in *in vivo* systems, where the interactions among cells and matrix environments are complex.

Therefore, simplified models that isolate microglia-EC interactions are needed to clarify their specific effects on endothelial barrier integrity and tight junction regulation.

In this study, we used multiscale *in vitro* models to dissect the role of microglia within NVU. We combined reductionist 2D co-cultures with a 3D perfused neuroimmune-vascular model incorporating human iPSC-derived microglia-like cells (iMGLs), neurons (iNs), astrocytes (iAs), and human brain microvascular endothelial cells (HBMECs). Across these platforms, iMGLs enhanced neuronal morphology and function, strengthened endothelial junctional organization, and preserved barrier integrity, including under AD-relevant inflammatory conditions. Together, these findings position microglia as key protective and regulatory components of the NVU, with direct implications for maintaining vascular integrity in neuroinflammation and neurodegeneration.

## 2. Results

### 2.1 Enhanced neuronal morphology and multicellular interactions in a 3D microfluidic neuroimmune-vascular model

In order to model the complex cellular interactions present in the NVU, we adapted an engineered microvessel platform we previously established to generate a neuroimmune-vascular model^61^. This model is comprised of iNs, iAs, and iMGLs in the collagen matrices, surrounding engineered microvessels lined with primary human brain microvascular endothelial cells (HBMECs) (NVU vessels) (Figure 1)^61, 62^. To build this model wild-type iMGLs were differentiated from hiPSCs using methods adapted from previously published protocol^63^. Concurrently, iNs, and iAs were differentiated from hiPSCs using a previously established neuronal differentiation protocol^64, 65^. iMGL were embedded within collagen hydrogels at 1×10^6^ cells/mL, alongside neurons and astrocytes at 3×10^6^ cells/mL (Figure 1A). The microvessel architecture consisted of two hexagonal networks connected by a straight channel, mimicking the branching nature of brain vasculature (Figure 1A). Individual vessel branches had an inner diameter of ∼100 µm, with open lumens that permitted continuous perfusion to achieve wall shear stress in the physiological range of brain capillaries despite the larger diameter. Engineered platforms were co-cultured for 7 days in optimized media (see Methods), allowing the ECs to form a continuous endothelium with luminal perfusion (Figure 1B orthogonal views), supporting neural cell survival, and enabling sustained 3D cell-cell interactions throughout the culture.

**Figure 1.**
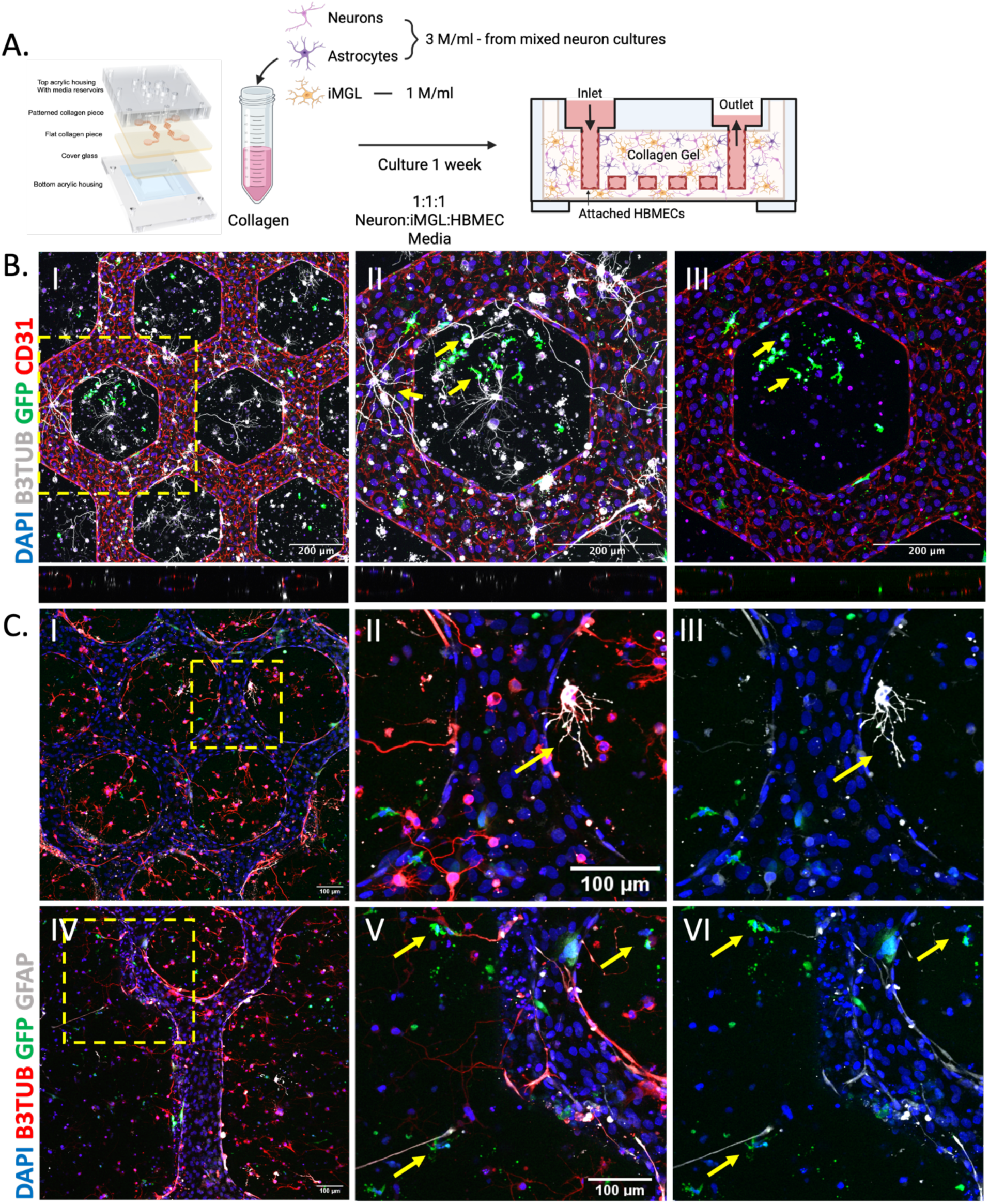
Development of a neuroimmune-vascular model. (A) Schematic overview of neuroimmune-vascular model fabrication. iPSC-derived neurons and astrocytes from mixed neuronal cultures were incorporated into a collagen gel at 3×10^6^ cells/ml at the time of collagen dilution and neutralization. Simultaneously, iMGL were incorporated into the collagen gel at a concentration of 1×10^6^ cells/ml. Vessels are then fabricated with a hexagon grid geometry and seeded with HBMECs. The hexagon grid pattern contains two hexagon structures connected by a straight channel. Engineered microvessels including these 3 cell types and primary HBMECs lining the vessel channels were fabricated (NVU vessels). Devices were fed a 1:1:1 media mixture of neuron:iMGL:HBMEC media for 7 days to establish a continuous endothelium and allow for iMGL-EC interactions. (B) Representative I)10x and II-III) 20x IF images of NVU vessels containing neurons (B3TUB, white), iMGL (GFP, green), and HBMECs (CD31, red). Scale bar = 100 μm. Yellow arrows cell-cell interactions of note between neurons and ECs and neurons and iMGL. Orthogonal views show the perfusable lumen of the NVU vessels. (C) Representative IF images of NVU vessels with neurons (B3TUB, red), iMGL (GFP, green), and astrocytes (GFAP, white). Scale bar = 100 μm. Arrows denote cell-cell interactions between astrocytes and vessels, iMGL and neurons, iMGL and astrocytes, and iMGL and neuronal cell debris.

We assessed the organization of neurons (βIII-tubulin, B3TUB), astrocytes (GFAP), endothelial cells (CD31), and GFP-labeled iMGL in our model (Figure 1B-C). Neurons displayed elongated processes with robust βIII-tubulin staining. Interactions between neurons and ECs, as well as neurons and iMGL, were also observed (Figure 1B arrows). Additionally, astrocytes extended GFAP+ end-feet toward the vessel surface, and iMGL closely associated with astrocytes and engaged with cellular debris (Figure 1C arrows). We compared neuronal morphology between complete NVU vessels and neuron-astrocyte vessels lacking iMGL (neuron vessels). NVU vessels exhibited stronger βIII-tubulin staining and significantly greater neurite outgrowth compared with vessels without iMGL (Supplemental Figure 1 A-C). Together, these results demonstrate the successful integration and dynamic interaction of all four NVU cell types within a single platform.

### 2.2 Inclusion of iMGL in neuron-astrocyte triculture promotes neural network connectivity and enhances neuron morphology

To understand individual cell-pair interactions, in particular, the role of microglia in neural network organization and function, we transitioned to a monolayer 2D triculture system consisting of iNs, iAs, and iMGLs. iMGLs were added to mixed neuronal cultures of iNs and iAs at a 1:1 ratio (Figure 2A). Immunocytochemistry (ICC) for microtubule associated protein 2 (MAP2) (iNs), glial fibrillary acidic protein (GFAP) (iAs), and ionized calcium-binding adaptor molecule 1 (IBA-1) (iMGLs) confirmed the presence of all three cell types in the triculture (Figure 2B).

**Figure 2.**
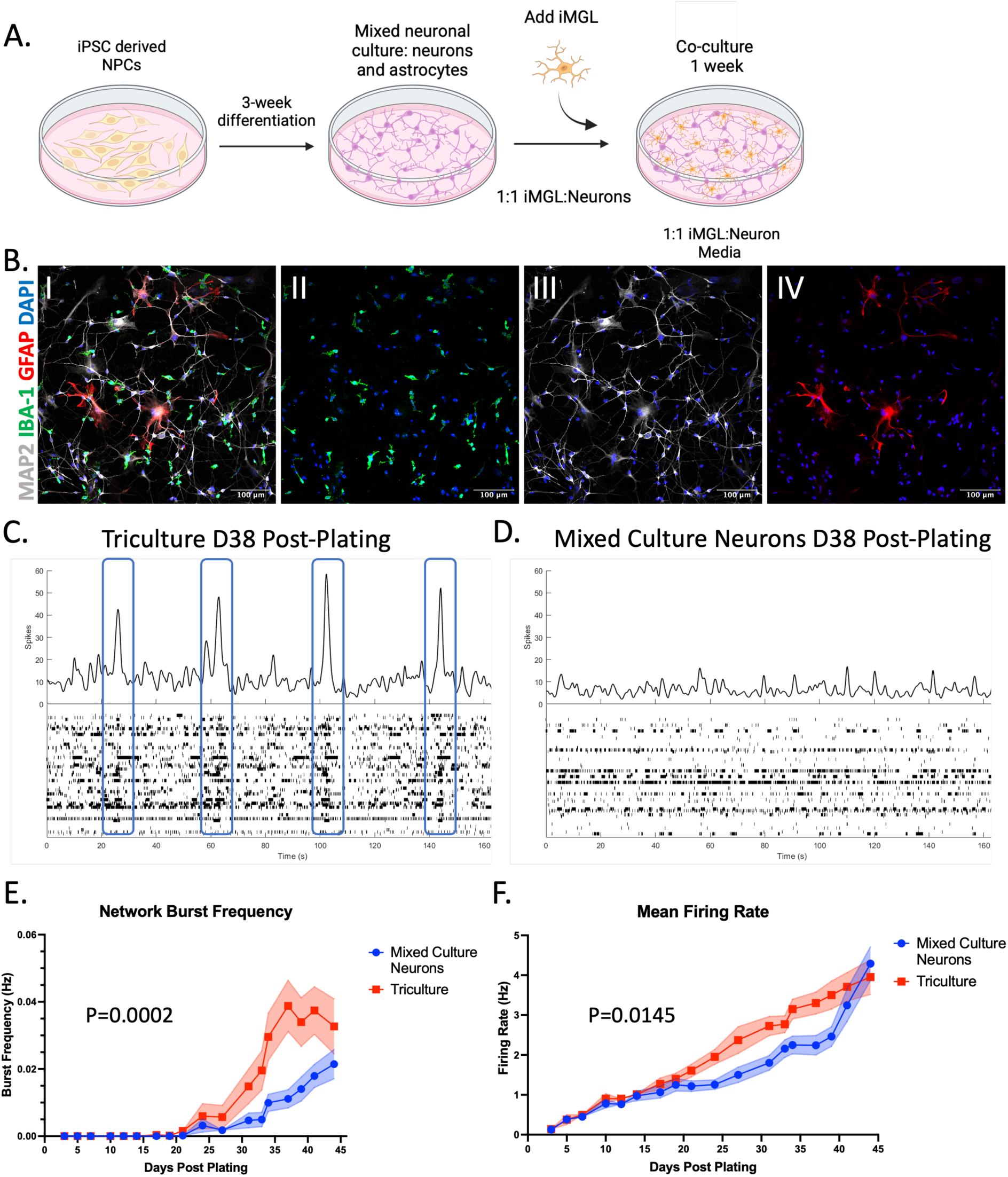
Tricultures show improved electrophysiological function compared to mixed culture neurons. (A) Schematic outlining the establishment of tricultures of neurons, astrocytes, and iMGL. iPSC-derived neural progenitor cells (NPCs) were first differentiated into mixed cultures of neurons and astrocytes for 3 weeks. iMGL were then added to the cultures at a 1:1 ratio with the total number of neurons and astrocytes and cultured in 1:1 neuron:iMGL media. Tricultures were maintained for 1 week. (B) Representative IF images of tricultures containing (II) iMGL (IBA-1), III) neurons (MAP2), IV) astrocytes (GFAP), and I) the composite of all cell types. Scale bar = 100 μm. IF images demonstrate our ability to co-culture these three key NVU cell-types concurrently. (C-D). Representative raster plots (bottom) and spike histograms (top) depicting the spiking activity of neurons in (C) tricultures or (D) mixed neurons and astrocytes cultured on a 6-well multielectrode array (MEA) plate for 38 days post-plating. Coordinated spiking across multiple electrodes results in peaks of activity in the spike histogram that are representative of network bursts and indicate coordinated activity across the culture (blue boxes). (E) Summary plot of the network burst frequency in tricultures vs. mixed culture neurons over 45 days. Quantification shows that there is a significantly higher network burst frequency in tricultures compared to mixed culture neurons. p=0.0002 (significance of the interaction of genotype and time) by two-way ANOVA. n=39-56 wells per culture condition across 4 biological replicates. (F) Summary plot of the mean firing rate in tricultures vs. mixed culture neurons over 45 days. Summary results demonstrate a significant increase in the overall electrical activity of tricultures compared to mixed culture neurons. p=0.0145 (significance of the interaction of genotype and time) by two-way ANOVA. n=74-75 wells per culture condition across 4 biological replicates.

We evaluated neural network electrical function using multielectrode array (MEA) recordings over 45 days. Overall activity was quantified via mean firing rate, while coordinated network connectivity was assessed using network burst frequency, defined as simultaneous spiking detected across multiple electrodes (Figure 2C, blue boxes). Over time, tricultures exhibited significantly higher network burst frequency and mean firing rate, reflecting enhanced synaptic coordination and overall electrical activity, compared to mixed neuron-astrocyte cultures without iMGLs (Figure 2C-F). Representative raster plots and spike histograms from day 38 post-plating further illustrate these differences in network bursting and electrical activity between tricultures and mixed neuronal cultures (Figure 2E,F).

Neurons in iMGL-containing tricultures exhibited stronger expression of the neuron markers, MAP2 and microtubule associated protein tau (MAPT), compared to neuron-astrocyte mixed cultures (Supplemental Figure 2A,C). These results suggest that iMGLs enhance neuronal health in line with recent work^66^. Neurite branch length, however, was not significantly altered by the presence of iMGLs in 2D (Supplemental Figure 2D). iMGLs in tricultures showed a more ramified phenotype, whereas iMGL monocultures displayed predominantly rod-like or amoeboid shapes (Supplemental Figure 2B), indicating a more activated state in monoculture. Collectively, these results suggest that the inclusion of iMGLs in hiPSC-derived brain tricultures promotes neuronal health and enhances coordinated neural network activity.

### 2.3 Co-culture of iMGL and HBMECs improves BBB-like structure of ECs and produces more ramified iMGLs

In addition to supporting neuronal health and network activity, microglia have been increasingly recognized as key modulators of brain vasculature^52–54, 67^. We therefore examined how microglia interact with brain ECs that constitute the BBB, a relatively understudied aspect of NVU biology.

To investigate the influence of microglia on EC junctional organization, we first established a simplified 2D EC-microglia co-culture model. Primary HBMECs were seeded to form a confluent monolayer for 48 h, after which iMGL were added (Figure 3A, C). Co-culture with iMGL (IBA-1⁺) increased the localization of the tight junction protein ZO-1 to the junctions of HBMECs compared with monocultures (Figure 3A I–III), which was confirmed by quantification of ZO-1 mean fluorescence intensity (MFI) at cell junctions (Figure 3B). Similarly, adherens junction protein, VE-Cadherin (VECAD), exhibited significantly greater junctional enrichment in EC-iMGL co-cultures compared to HBMEC monocultures (Figure 3C I–III, 3D). Together, these findings indicate that iMGLs enhance the junctional localization of both tight and adherens junction proteins, potentially promoting a more BBB-like endothelial phenotype in 2D co-culture.

**Figure 3.**
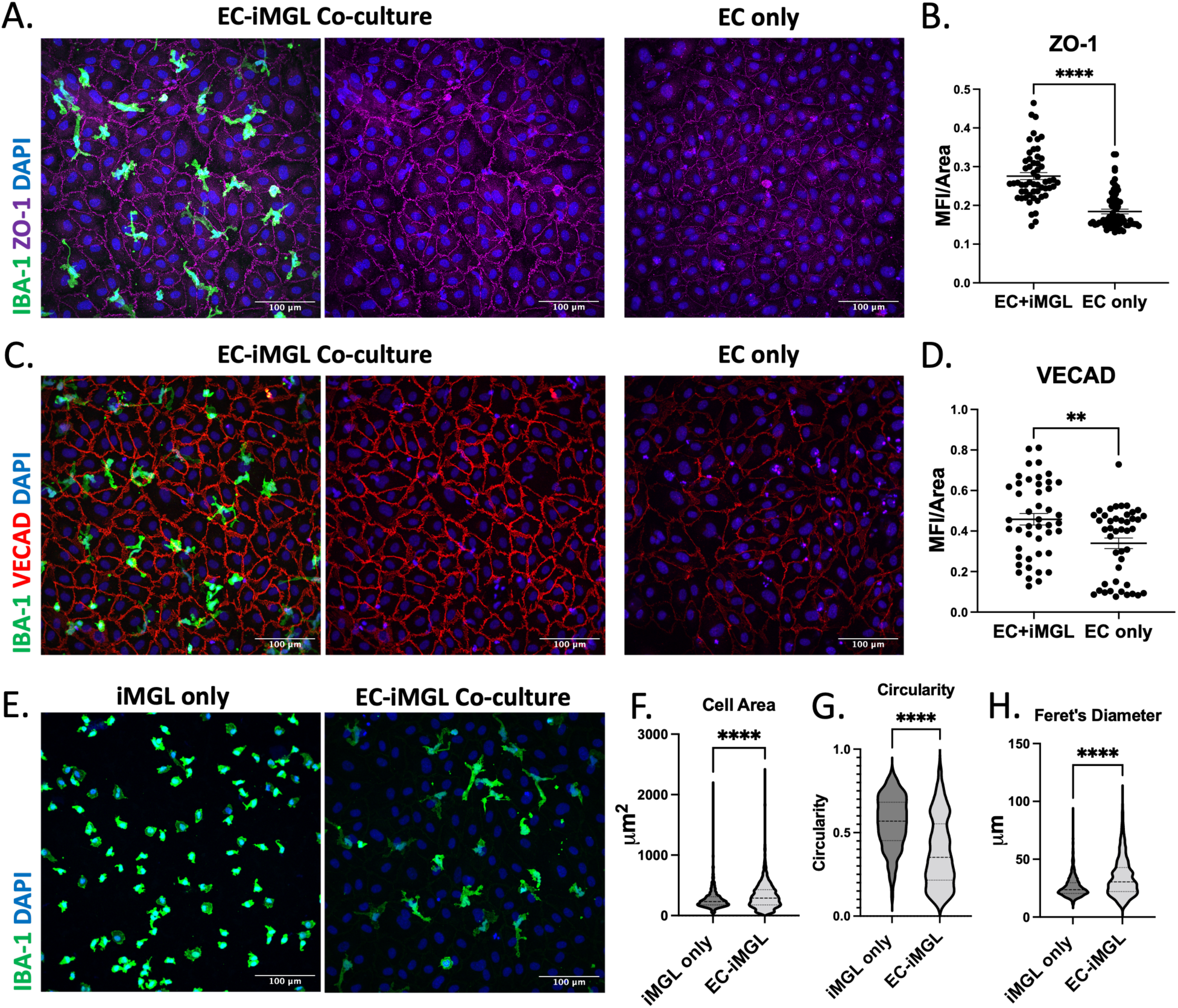
iMGL co-culture upregulates endothelial tight and adherens junction protein localization and promotes a ramified microglial morphology. (A) Representative immunofluorescence (IF) images of human brain microvascular endothelial cells (HBMECs) stained for ZO-1 (magenta) and cultured (I-II) with iMGL (IBA-1, green) or (III) in monoculture. Scale bar = 100 μm. (B) Quantification of ZO-1 mean fluorescence intensity (MFI) normalized to ZO-1 positive area. n = 3 biological replicates. (C) Representative IF images of HBMECs stained for VE-Cadherin (VECAD, red) and cultured (I-II) with iMGL (IBA-1, green) or (III) in monoculture. Scale bar = 100 μm. (D) Quantification of VECAD MFI normalized to VECAD positive area. n = 3 biological replicates. (E) Representative IF images of iMGL stained for IBA-1 (green) and cultured (I) in monoculture or (II) with HBMECs (EC-iMGL). Scale bar = 100 μm. (F) Quantification of cell area in µm^2^ of individual iMGL in cultures of iMGL only vs. iMGL co-cultured with HBMECs (EC-iMGL). iMGL in EC-iMGL co-cultures have a larger size indicated by an increase in cell area. n = 4 biological replicates. (G) Quantification of cell circularity (4π (area/perimeter^2^)) of individual iMGL in cultures of iMGL only vs. iMGL co-cultured with HBMECs (EC-iMGL). Circularity values of 1.0 correspond to circular cells, whereas values near zero correspond to elongated or ramified microglia. iMGL in monocultures have higher circularity than EC-iMGL co-cultures. n = 4 biological replicates. (H) Quantification of Feret’s diameter in µm of individual iMGL in cultures of iMGL only vs. iMGL co-cultured with HBMECs (EC-iMGL). Feret’s diameter represents the longest distance between two points along the perimeter of the cell. Co-cultured iMGL have a longer Feret’s diameter compared to monocultured iMGL. n = 4 biological replicates. All plots: Error bars, mean ± SEM. *p < 0.05, **p < 0.002, ***p < 0.0002, ****p<0.0001 by unpaired t-test.

We next assessed how co-culture influenced iMGL morphology, an indicator of activation state^68^. Resting microglia typically exhibit a branched, ramified morphology suited for environmental surveillance, whereas activated microglia adopt an amoeboid, unbranched form to facilitate motility^68^. Here, iMGLs in co-culture with ECs displayed more complex, ramified morphologies (Figure 3E II), whereas monocultured iMGL were predominantly amoeboid (Figure 3E I). Morphological analysis using established parameters^69^ confirmed these observations. Co-cultured iMGLs had significantly greater cell area (Figure 3F), lower circularity (Figure 3G), and longer Feret’s diameter (Figure 3H) than monocultures. These changes are consistent with a shift toward a less activated microglial state in the presence of endothelial cells.

### 2.4 HBMEC treatment with iMGL conditioned media (CM) partially recapitulates increases in junctional protein expression observed in EC-iMGL co-cultures

To understand the mechanism behind the increased tight and adherens junction localization in EC-iMGL co-cultures, we next investigated whether these effects were mediated by soluble factors secreted by iMGL or required direct cell-cell contact. Conditioned media (CM) from iMGL cultures were applied to HBMEC monocultures, and junctional protein localization and expression was compared across CM-treated HBMECs, HBMEC monocultures, and direct EC-iMGL co-cultures. This approach enabled us to distinguish contact-dependent from secreted factor-mediated effects.

We performed a western blot to evaluate total VECAD protein levels in CM-treated HBMECs, HBMEC monocultures, and direct EC-iMGL co-cultures and saw no differences in overall protein expression between conditions (Supplemental Figure 3). However, ICC for VECAD (Figure 4B) displayed a graded pattern where monocultures had the lowest MFI, CM-treated cultures showed intermediate MFI, and direct co-cultures had the highest MFI (Figure 4D). This suggests that differences in MFI were representative of changes in protein localization as opposed to changes in overall protein expression. Visualization and quantification of ZO-1 MFI demonstrated that CM-treated HBMECs and direct co-cultures exhibited comparable ZO-1 localization, both significantly higher than monocultures (Figure 4A,C). Therefore, while conditioned media from iMGL was sufficient to fully restore ZO-1 levels in HBMECs, VECAD junctional protein localization showed only a partial increase compared with direct co-culture. This distinction suggests that tight junction regulation may be largely driven by soluble microglial factors, whereas adherens junction enhancement requires additional signals provided by direct microglia-endothelial contact.

**Figure 4.**
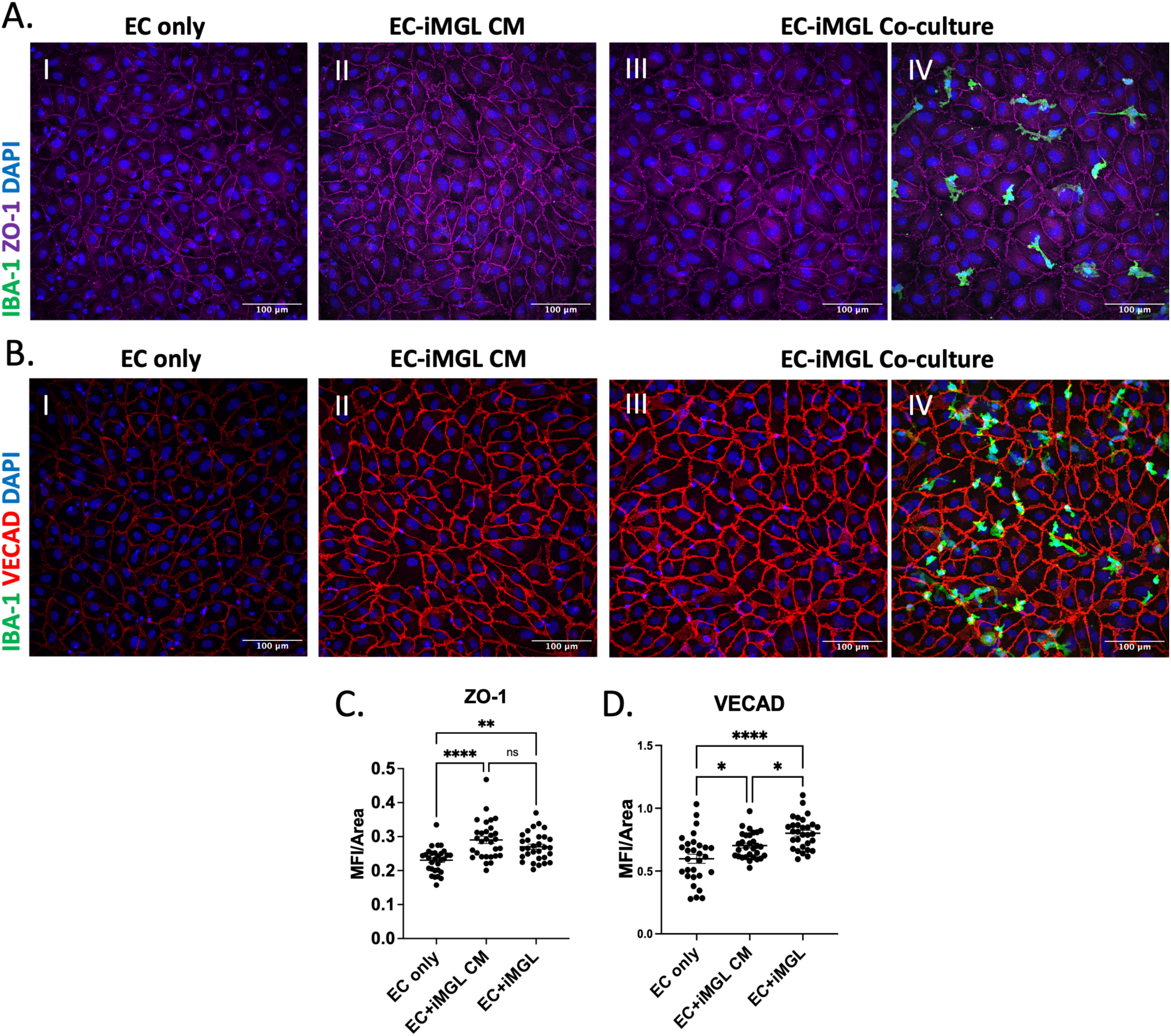
iMGL-conditioned media elevates EC junction protein expression. (A) Representative IF images of HBMECs stained for ZO-1 (magenta) and cultured (I) in monoculture (II) with iMGL conditioned media (CM), or (III-IV) with iMGL (IBA-1, green). Scale bar = 100 μm. (B) Representative IF images of HBMECs stained for VE-Cadherin (VECAD, red) and cultured (I) in monoculture (II) with iMGL conditioned media, or (III-IV) with iMGL (IBA-1, green). Scale bar = 100 μm. (C) Quantification of ZO-1 MFI normalized to ZO-1 positive area in cultures of HBMECs only, HBMECs treated with iMGL CM, or HBMECs cultured directly with iMGL (EC-iMGL). Direct co-cultures and CM treated cultures had higher ZO-1 MFI than EC only cultures but were not significantly different from each other. n = 3 biological replicates. (D) Quantification of VECAD MFI normalized to VECAD positive area in cultures of HBMECs only, HBMECs treated with iMGL CM, or HBMECs cultured directly with iMGL (EC-iMGL). EC only cultures had lowest VECAD MFI, CM treated cultures were intermediate, and direct co-cultures had the highest MFI. n = 3 biological replicates. All plots: Error bars, mean ± SEM. *p < 0.05, **p < 0.002, ***p < 0.0002, ****p<0.0001 by one-way ANOVA with Tukey’s correction for multiple comparisons test.

### 2.5 In co-cultures, iMGL attenuate the inflammatory response of HBMEC following TNFα stimulation

To determine how iMGLs influence the EC response to inflammatory stimulation in our co-culture model, we focused on TNFα, an AD-relevant cytokine. TNFα levels are elevated in both the serum and cerebrospinal fluid (CSF) of patients with AD and higher baseline levels of TNFα correlate with a greater degree of cognitive decline^70–75^. We therefore used TNFα to perturb our model and assessed the EC inflammatory response by visualizing intercellular adhesion molecule 1 (ICAM1) expression (Figure 5A,B).

**Figure 5.**
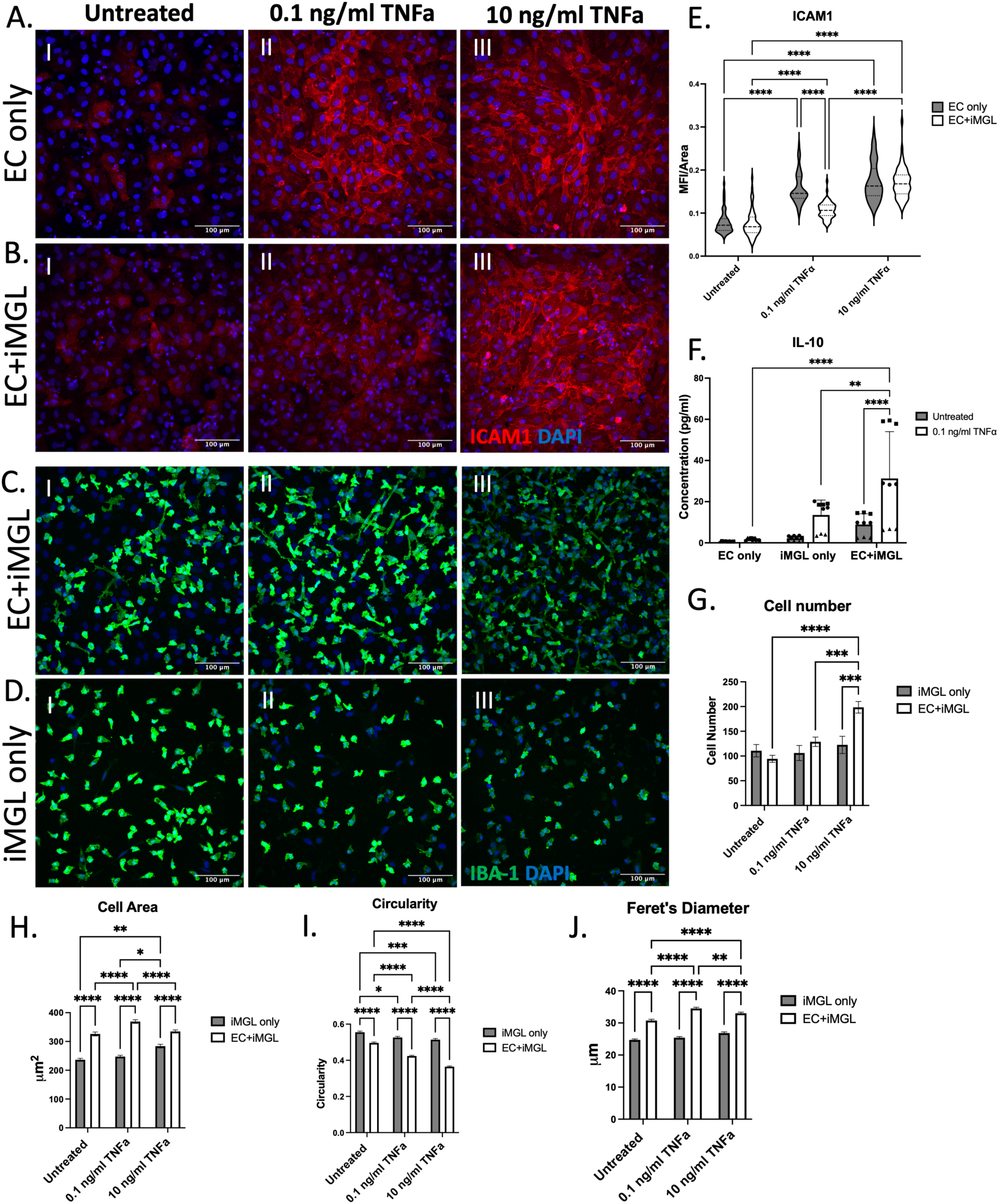
iMGLs attenuate the inflammatory response of HBMECs after 18 h stimulation with 0.1 ng/mL TNFα. (A-B) Representative IF images of HBMECs (A) in monoculture (EC only) or (B) co-cultured with iMGL (EC+iMGL) I) untreated, II) treated with 0.1 ng/ml TNF⍺, or III) treated with 10 ng/ml TNF⍺ for 18 hours. Staining for intercellular adhesion molecule 1 (ICAM1, red). Scale bar = 100 μm. (C-D) Representative IF images of iMGL (IBA-1, green) in (C) EC+iMGL co-cultures or (D) in monocultures I) untreated, II) treated with 0.1 ng/ml TNF⍺, or III) treated with 10 ng/ml TNF⍺ for 18 hours. Scale bar = 100 μm. (E) Quantification of ICAM1 MFI normalized to ICAM1 positive area (MFI/Area) in EC only or EC+iMGL cultures untreated or treated with TNF⍺. n = 3 biological replicates. Two-way ANOVA with Šídák’s multiple comparisons test. (F) Secreted IL-10 levels (pg/ml) in the media of EC only, iMGL only, or EC-iMGL cultures as measured by Meso Scale Discovery (MSD) immunoassay with or without 0.1 ng/ml TNF⍺ treatment. n=3 biological replicates. Mixed-effects analysis with Tukey’s multiple comparisons test. (G) Quantification of iMGL cell number per image in EC only or EC+iMGL cultures untreated or treated with TNF⍺. (H) Quantification of cell area in µm^2^. n = 3 biological replicates. (I) Quantification of cell circularity (4π (area/perimeter^2^)). Circularity values of 1.0 correspond to circular cells, whereas values near zero correspond to elongated or ramified microglia. n = 3 biological replicates. (J) Quantification of Feret’s diameter in µm. Feret’s diameter represents the longest distance between two points along the perimeter of the cell. n = 3 biological replicates. (G-J) Two-way ANOVA with Tukey’s multiple comparisons test. All plots: Error bars, mean ± SEM. *p < 0.05, **p < 0.002, ***p < 0.0002, ****p<0.0001.

Monocultures of HBMECs and HBMEC-iMGL co-cultures were treated with no TNFα, moderate (0.1 ng/ml) TNFα, or high (10 ng/ml) TNFα for 18 hours. ICAM1 expression was comparable between EC monocultures and co-cultures under both untreated and high TNFα conditions (Figure 5A,B, panels I and III). In contrast, moderate TNFα treatment resulted in significantly lower ICAM1 expression in co-cultures compared with EC monocultures (Figure 5A,B, panel II), a finding confirmed by quantification of ICAM1 MFI (Figure 5E). This suggests that iMGL may help blunt the EC inflammatory response in co-cultures after acute inflammatory stimulation.

When visualizing iMGL number and morphology in these cultures, we observed a significant increase in the number of iMGL per field of view in EC-iMGL co-cultures treated with increasing doses of TNFα (Figure 5C,F). In contrast, iMGL number remained unchanged in monocultures (Figure 5D,F). This difference may be driven by TNFα-induced upregulation of endothelial adhesion molecules such as ICAM1, which could promote iMGL attachment within co-cultures.

Previous studies have reported that TNFα stimulation can induce a more elongated microglial morphology, in contrast to the amoeboid shape typically associated with activated microglia^76^. Consistent with this, we observed an increase in cell area in iMGL monocultures and a decrease in circularity in both monocultures and EC-iMGL co-cultures upon TNFα stimulation (Figure 5H,I). More rod-like iMGL were also apparent in TNFα-treated co-cultures (Figure 5C). Surprisingly, however, Feret’s diameter did not significantly change in either culture condition with increasing TNFα (Figure 5J).

To complement these findings, we quantified pro-and anti-inflammatory cytokine levels in culture media using a Mesoscale Discovery (MSD) assay. Media from EC monocultures, iMGL monocultures, and EC-iMGL co-cultures, with and without TNFα stimulation, were analyzed for IL-10, IL-4, IL-13, IFN-γ, IL-12p70, IL-1β, IL-2, and IL-6 (Supplemental Figures 4,5). Interestingly, for pro-inflammatory cytokines (IFN-γ, IL-12p70, IL-1β, IL-2, and IL-6), TNFα-stimulated co-cultures consistently showed lower levels than would be expected from the sum of monocultures (Supplemental Figure 5). These findings suggest that ECs, iMGL, or their interactions suppress inflammatory cytokine output, resulting in reduced activation in the co-culture context compared to monoculture. Additionally, under moderate TNFα stimulation, co-cultures secreted significantly higher levels of the anti-inflammatory cytokine IL-10 compared with monocultures (Figure 5F). This elevated IL-10 release may underlie the dampened endothelial inflammatory response observed in our co-cultures.

### 2.6 Barrier integrity analysis in a 3D neuroimmune-vascular model indicates that iMGLs contribute to maintaining EC barrier function

To evaluate whether iMGL influence vessel barrier function, we developed a reductionist 3D model by incorporating only iMGLs in the matrices around the endothelium. The advantages of this system are that it allows for the proper spatial arrangement of iMGL relative to ECs, contains perfusable microvessels, and has both branched vascular architecture as well as a straight channels suitable for functional permeability measurements.

We fabricated perfusable microvessels and included iMGL at 1×10^6^ cells/mL in the bulk collagen gel used for fabrication. HBMECs were, again, used to seed the engineered microvessels (Figure 6A,B). ICC of these 3D iMGL-vessel cultures revealed that a subset of iMGLs adhered to the vessel surface, while others remained distributed within the surrounding collagen matrix (Figure 6B). When these vessels lumens were perfused with 70 kDa FITC-dextran, diffusion across the EC barrier was imaged over 5 minutes. Barrier function was quantified using a permeation index, calculated as the normalized dextran permeation distance over 2 or 5 minutes. Compared with EC-only vessels maintained in identical media, iMGL vessels displayed an approximately 2.3-fold lower permeation index (Figure 6C-E), indicating that iMGL presence enhances EC barrier integrity. These findings indicate that iMGL not only interact structurally with vessels but also actively support barrier function.

**Figure 6.**
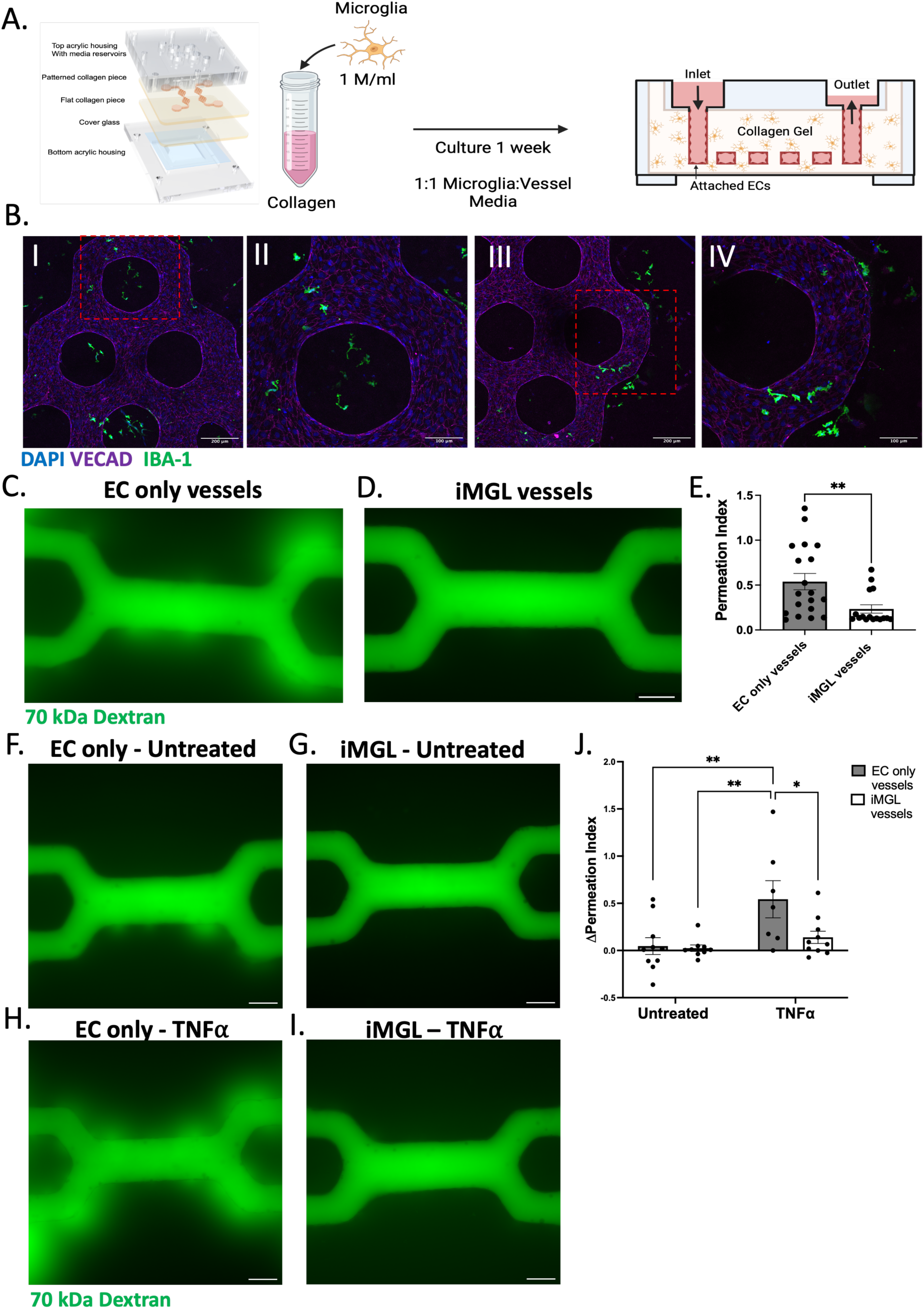
iMGL co-culture with engineered microvessels stabilizes the endothelial barrier under basal conditions. (A) Schematic of iMGL containing engineered microvessel fabrication (iMGL vessels). iMGL are mixed into the collagen gel at a density of 1 million cells/ml. iMGL or EC only vessels were cultured under gravity driven flow using 1:1 iMGL:HBMEC media for 1 week. (B) Representative IF images of iMGL vessels stained for VECAD (magenta) and IBA-1 (green). iMGL interact with the EC surface in engineered microvessels. Scale bar = 200 μm (I and III) or 100 μm (II and IV). (C-D) Representative images of (C) EC only or (D) iMGL vessels perfused with 70 kDa FITC-dextran to measure the permeability of the vessel barrier. (D) Scale bar = 150 μm. (E) Quantification of the permeation index of EC only or iMGL vessels following 70 kDa FITC dextran perfusion for 5 mins. Permeation index is a dimensionless parameter that represents the normalized distance traveled by dextran over the course of 5 mins. Error bars, mean ± SEM. *p < 0.05, **p < 0.002, ***p < 0.0002, ****p<0.0001 by unpaired t-test. (F-G) Representative images of untreated (F) EC only or (G) iMGL vessels perfused with 70 kDa FITC-dextran. Scale bar = 150 μm. (H-I) Representative images of 18-hour 0.1 ng/ml TNF⍺ treated (H) EC only or (I) iMGL vessels perfused with 70 kDa FITC-dextran. Scale bar = 150 μm. (J) Quantification of the change in permeation index of EC only or iMGL vessels to 70 kDa FITC dextran after 2 mins of dextran perfusion. Change in permeation index was calculated by subtracting the permeation index before TNF⍺ treatment (baseline) from the permeation index after treatment with TNF⍺ (treatment). n=7-10 vessels per condition across 3 biological replicates. Error bars, mean ± SEM. *p < 0.05, **p < 0.002, ***p < 0.0002, ****p<0.0001 by two-way ANOVA with Šídák’s multiple comparisons test.

### 2.7 iMGL prevent EC barrier breakdown following TNFα stimulation

We next investigated whether microglia exert a protective or detrimental effect on endothelial barrier function following barrier-disruptive stimulation. TNFα, a known inducer of BBB dysfunction^77, 78^, was used at 0.1 ng/ml for 18 hours. Barrier integrity was assessed by measuring the permeation index of vessels before and after treatment.

In EC-only vessels, TNFα markedly increased dextran diffusion across the barrier compared with untreated controls (Figure 6F,H). In contrast, iMGL vessels exposed to TNFα exhibited minimal dextran leakage, with permeability only slightly elevated relative to untreated controls (Figure 6G,I). Quantification of the change in permeation index confirmed that untreated EC-only and iMGL vessels showed near-zero mean changes (0.0466 and 0.0243, respectively) (Figure 6J). TNFα-treated EC-only vessels displayed a significantly higher positive change in permeation index, indicating substantial barrier disruption (Figure 6J). Notably, although TNFα-treated iMGL vessels showed a slight increase compared with untreated controls, the change was 3.9-fold lower than in EC-only vessels (0.1395 vs. 0.5427) (Figure 6J). These findings indicate that in response to acute TNFα challenge, iMGL substantially attenuate EC barrier breakdown, supporting a protective role for microglia under short-term inflammatory conditions.

## 3. Discussion

NVU dysfunction is a well-established feature of AD, yet the specific contributions of microglia remain poorly understood. Prior studies examining EC-microglia interactions in healthy and disease contexts have been limited, with contradictory results ^54–57, 60^. Here, we addressed this gap using a series of human *in vitro* NVU models of increasing complexity to enable direct interrogation of microglial influence on endothelial and neuronal function. We found that co-culture with iMGL upregulated tight and adherens junction protein localization in HBMECs, an effect partially mediated by paracrine signaling. In 3D engineered microvessels, iMGL enhanced barrier integrity. Following moderate levels of TNFα exposure, iMGL dampened endothelial inflammatory activation in 2D and prevented barrier breakdown in 3D. Extending our study to multicellular NVU models, we observed that iMGL promoted neuronal morphology and network function. Together, these findings provide direct evidence that microglia exert protective and regulatory roles within the NVU, supporting both BBB stability and neuronal health, and underscore their importance in maintaining NVU function in health and disease contexts.

We established a 3D neuroimmune-vascular model that integrates primary HBMECs with iPSC-derived microglia-like cells, neurons, and astrocytes in a single culture system. While several microfluidic NVU/BBB platforms have incorporated microglia, these models rarely dissect microglia-endothelial interactions or directly evaluate the role of microglia within the NVU^45, 50, 51, 79^. By integrating simplified EC-iMGL co-cultures with multicellular NVU constructs, our approach directly addresses this gap and provides a framework to dissect how microglia regulate BBB integrity and neuronal health. This is especially important given that recent transcriptomic and proteomic studies highlight fundamental differences between human and mouse microglia, particularly in AD-relevant pathways^80–82^. Thus, our models not only reveal microglial contributions to NVU function but also offer a human-specific platform for mechanistic discovery and translational applications in neurodegenerative disease research.

We observed increased neuronal activity, reflected by higher mean firing rates, and enhanced network connectivity, reflected by elevated network burst frequency, in neuronal cultures containing iMGL. The increase in network burst frequency indicates that there are stronger synaptic connections between neurons when cultured with iMGL^83^. This is consistent with known microglial functions, including secretion of anti-inflammatory cytokines and trophic factors that support neuronal health, as well as activity-dependent synaptic pruning that strengthens active circuits^84, 85^. Our results align with recent findings that microglia co-culture with neurons *in vitro* leads to significant increases in overall electrical activity and a trend toward increased network bursting^66^. In addition, we observed morphological changes in neurons in 2D culture with iMGL and our 3D neuroimmune-vascular model containing iMGL. Interestingly, differences in neurite extension were more pronounced in the 3D NVU model than in 2D cultures. Previous work has shown that neurons in 3D hydrogels display greater process outgrowth^86, 87^, underscoring the advantages of 3D culture systems in recapitulating *in vivo*-like neuronal architecture.

We found that culturing HBMECs with iMGL-conditioned media partially reproduced the upregulation of junctional proteins observed in direct co-culture. ZO-1 junctional protein localization was comparable between direct co-cultures and CM-treated HBMECs, whereas VECAD localization was elevated in CM-treated cultures but did not reach the levels observed with direct contact. These findings align with prior reports that microglial secreted factors, including trophic molecules such as Insulin-Like Growth Factor-1 (IGF-1), can enhance endothelial junctional proteins and support BBB function^88–91^. In addition, paracrine CSF-1R signaling between microglia and brain ECs has been implicated in maintaining tight junction expression^59^. At the same time, microglia have also been shown to physically interact with ECs, for example by delivering Claudin-5 following inflammatory challenge, thereby directly reinforcing barrier integrity^54^. Our data suggest that both soluble and contact-dependent mechanisms contribute to microglial support of endothelial junctional integrity. The differential effects on ZO-1 and VECAD further raise the possibility that distinct regulatory pathways underlie tight versus adherens junction modulation by microglia, an important question for future investigation.

In our 3D perfused microvessel model, the presence of iMGL supported endothelial barrier integrity. This observation contrasts with a prior *in vivo* study reporting that one month of microglial depletion did not disrupt BBB structure or function^4^. One possible explanation for this difference is that other NVU cell types, such as astrocytes or pericytes, may compensate for the chronic absence of microglia *in vivo*, thereby masking their contribution. Alternatively, microglia may exert a modest but meaningful influence on BBB maintenance that becomes more apparent in reductionist *in vitro* systems, where the dominant contributions of astrocytes and other support cells are absent^92^. These possibilities highlight the importance of context in assessing microglial roles at the BBB and suggest that microglial contributions, while potentially subtle in intact tissue, may be critical under specific physiological or pathological conditions.

Upon TNFα stimulation (0.1 ng/ml; 100 pg/ml), we observed reduced ICAM1 upregulation in iMGL-HBMEC co-cultures compared to EC monocultures. Notably, this concentration is close to levels reported in the serum of AD patients (∼73 pg/ml)^93^, suggesting that microglia can attenuate endothelial inflammatory activation under physiologically relevant conditions. In parallel, we detected increased secretion of the anti-inflammatory cytokine IL-10 in TNFα-treated co-cultures relative to monocultures. Given that IL-10 has been shown to directly suppress ICAM1 expression on ECs^94, 95^, enhanced IL-10 production in co-cultures may represent one mechanism by which microglia dampen EC inflammatory responses.

Importantly, iMGLs also prevented barrier breakdown following acute TNFα exposure. Consistent with our findings, IL-10 has previously been shown to preserve barrier integrity *in vitro* by inhibiting EC apoptosis and tight junction loss^96^. This supports a role for microglial secreted factors in BBB protection. Beyond paracrine signaling, microglial physical interactions with ECs might also be critical in maintaining BBB integrity.

While the 2D and 3D models developed here proved useful for investigating microglial interactions with neurons and ECs, they are not without limitations. In the 2D system, iMGL interact with ECs on the apical (luminal) surface, whereas *in vivo*, microglia primarily contact brain ECs from the basolateral (abluminal) side^97^. We addressed this by developing 3D microvessels with abluminally positioned iMGLs. Additional refinements could further increase physiological relevance of our model, such as incorporating more brain-specific extracellular matrix proteins (e.g., collagen type III and IV, fibronectin, vitronectin)^98^. Moreover, our focus was on microglial responses to neuroinflammatory stimuli, rather than interactions with Aβ or tau, classical hallmarks of AD. Notably, inflammation may precede protein aggregation in AD^99^, so the findings presented here may be indicative of early pathogenic events. Finally, future studies should extend TNFα exposure from 18 hours to several days to test whether iMGL undergo a functional switch toward detrimental effects on BBB integrity, as previously reported^54^.

## 4. Conclusion

In this study, we used multiscale *in vitro* models to investigate the role of microglia within the NVU. Employing models ranging from 2D co-cultures to a 3D perfused neuroimmune-vascular system composed of human iMGLs, iNs, iAs, and HBMECs, we found that iMGL enhanced neuronal structure and function, improved endothelial junctional organization, and helped maintain barrier integrity under AD-related inflammatory conditions. These findings position microglia as active regulators of NVU homeostasis and support their potential role in preserving vascular integrity during neuroinflammatory and neurodegenerative states.

## 5. Methods

### iPSC-derived microglia-like cell (iMGL) differentiation

Induced microglia-like cells (iMGLs) were generated from human induced pluripotent stem cells (hiPSCs) using a modified version of the protocol described by McQuade et al. 2018, with adaptations outlined in Mishra et al. 2025^63, 100^. Wild-type (WT) iMGLs were derived from a previously characterized control hiPSC line^64^ from a male donor with an APOE ε3/ε4 genotype^101^.

hiPSCs were maintained in mTeSR Plus medium supplemented with 10 µM ROCK inhibitor Y-27632 (#A3008; ApexBio) on 6-well plates (#657160; CELLSTAR) coated with 1:30 growth factor-reduced Matrigel (#356231; Corning). Cells were fed daily until they reached ∼70% confluency.

To initiate hematopoietic differentiation, hiPSCs were lifted using ReLeSR (#100-0483; STEMCELL Technologies) and replated as aggregates (∼40 cells/aggregate, 20–40 aggregates/well) in mTeSR Plus + ROCK inhibitor on 1:30 Matrigel-coated plates.

Approximately 10 colonies attached per well, which was optimal for hematopoietic progenitor cell (HPC) generation. On day 0, medium was switched to STEMdiff Hematopoietic Supplement A (#05310; STEMCELL Technologies), followed by a medium change to Supplement B on day 3 upon embryoid body formation. Cultures were maintained in Supplement B for 7 days with partial media changes every other day until HPCs were produced from embryoid bodies. At this point, media was added but not removed from the cultures to preserve HPCs.

On day 12, non-adherent HPCs were harvested by collecting the supernatant and performing gentle PBS washes. HPCs were cryopreserved in Bambanker cell freezing medium (#BBH01; Bulldog-Bio) at 2 × 10⁶ cells/vial.

For microglial differentiation, thawed HPCs were plated at 2 × 10⁵ live cells/well on 1:60 Matrigel-coated 6-well plates in microglia differentiation medium composed of DMEM/F12 without phenol red (#11039047; Thermo Fisher Scientific), supplemented with Insulin-Transferrin-Selenium-Sodium Pyruvate (#51300044), B27 (#17504044), N2 (#17502048), GlutaMAX (#35050061), nonessential amino acids (#11140050), monothioglycerol (#M1753; Sigma), and insulin (#I2643; Sigma). Freshly added cytokines included TGF-β (#130-108-969; Miltenyi Biotec), IL-34 (#200-34; Peprotech), and M-CSF (#PHC9501; Thermo Fisher Scientific). Cells were fed every other day for 24 days.

On day 24, CD200 (#C311; Novoprotein) and CX3CL1 (#300-31; Peprotech) were added to the medium to promote microglial maturation. After 3 days, cells were returned to the standard differentiation medium and maintained until day 32. Differentiated iMGLs were either used directly in experiments or cryopreserved at 2–3×10⁶ cells/vial in Bambanker freezing medium (#BB05; Bulldog-Bio).

### Human brain microvascular cell (HBMEC) Culture

HBMECs (#ACBRI 376 V; Cell systems) from passage number 3–6 were used for experimentation. HBMECs were maintained in T75 cell culture flasks and fed HBMEC growth medium (GM) (#3202; EGM-2 MV Microvascular Endothelial Cell Growth Medium-2 BulletKit (#3156; EBM-2 Basal Medium), 5% fetal bovine serum (FBS), hydrocortisone, human fibroblast growth factor-beta (hFGF-B), vascular endothelial growth factor (VEGF), R3-insulin-like growth factor-1 (R3-IGF-1), ascorbic acid, human epidermal growth factor (hEGF), GA-1000 (gentamycin and amphotericin)). Cells were maintained at 37 °C in a 5% CO2 incubator and media was replaced every 2-3 days.

### iMGL-HBMEC 2D co-cultures

Human brain microvascular endothelial cells (HBMECs) were cultured as described above. Upon reaching confluence, cells were passaged using 0.05% Trypsin-EDTA (#25300054; Gibco) and seeded onto 1:100 Geltrex-coated Ibidi µ-Slide 8-well chamber slides (#80806; Ibidi) at a density of 25,000 cells/well in HBMEC growth medium (GM). HBMECs were maintained for 48 hours to allow formation of a confluent monolayer.

Following monolayer formation, the medium was replaced with a 1:1 mixture of HBMEC GM and microglia differentiation medium. WT iMGLs were then added at a density of 200,000 cells/well directly onto the HBMEC monolayer. Control conditions included (1) HBMEC-only wells without iMGLs and (2) iMGL-only wells without HBMECs, in which iMGLs were seeded at the same density at the time of co-culture initiation.

All experimental conditions - HBMEC-only, iMGL-only, and HBMEC–iMGL co-culture - were maintained in the same 1:1 mixture of HBMEC GM and microglia differentiation medium for the duration of the culture. Media were gently added (without aspiration, to prevent loss of semi-adherent iMGLs) on alternate days. Cultures were maintained for 5 days.

After 5 days, cultures were either fixed in 4% paraformaldehyde (PFA; #J19943.K2; Thermo Fisher Scientific) for immunocytochemistry or treated with tumor necrosis factor-alpha (TNFα) as described below. Co-culture experiments were performed with three biological replicates, each consisting of HBMECs from independent passages and iMGLs from separate differentiation batches. For each biological replicate, two technical replicates were plated in separate wells of an 8-well Ibidi chamber slide.

### iMGL conditioned medium (CM) collection and application

For each conditioned media (CM) replicate, iMGL were thawed and plated in a single well of a 6-well plate coated with 1:60 Matrigel at a density of 2×10^6^ cells/well. CM was collected from two independent WT clones. Cells were allowed to recover for three days post-thaw. On day 3, the medium was replaced with 2 mL of a 1:1 mixture of HBMEC growth medium and microglia differentiation medium (supplemented with mCSF, TGF-β, and IL-34). For the following six days, 0.5 mL of fresh 1:1 medium was added to the cultures daily. After seven days of conditioning, the media were collected and centrifuged to remove any floating cells. The CM was then aliquoted and stored at –80 °C until further use.

To compare the effect of CM with direct co-cultures, 2D co-cultures were generated as described above. A set of HBMEC only control wells was also included. In the CM treated condition, HBMECs were allowed to form a monolayer as described above. At the same time iMGL were added to co-cultures, 200 μL CM supplemented with mCSF, TGF-β, and IL-34 was applied to wells of HBMECs. 100 μL of CM was added to the wells on alternating days for 5 days. Following 5 days of culture, CM treated wells, EC-iMGL co-cultures, and EC only wells were either fixed for ICC or lysates were collected for western blot.

### TNF**α** treatment of 2D co-cultures

Tumor necrosis factor-α (TNFα) (#H8916, Sigma-Aldrich) was reconstituted in 0.1% BSA (A7906; Sigma-Aldrich) in PBS at a concentration of 10 µg/mL. Cultures of iMGL-HBMEC co-cultures, HBMEC only, and iMGL only were generated as described above. These cultures were treated with TNFα as follows: All but 100 µL of media was carefully removed from each well to minimize the loss of microglia in the iMGL-only and iMGL-HBMEC co-culture conditions, due to the semi-adherent nature of the microglia.

The 10 µg/mL TNFα stock was diluted in a 1:1 mixture of HBMEC GM and microglia differentiation medium (referred to as 1:1 medium) to obtain working concentrations of 0.15 ng/mL or 15 ng/mL. Then, 200 µL of either TNFα dilution was added to the appropriate wells, bringing the final media volume in each well to 300 µL, and resulting in final TNFα concentrations of 0.1 ng/mL or 10 ng/mL.

For untreated control wells, 200 µL of 1:1 medium without TNFα was added to reach the same final volume and maintain consistency in media composition across all conditions. All cultures, iMGL-HBMEC co-cultures, HBMEC-only, and iMGL-only, were incubated with either untreated or TNFα-containing media for 18 hours. At the end of this incubation, the media from these cultures were collected for cytokine analysis.

### iMGL morphology quantification

iMGL morphology was quantified using custom ImageJ scripts. IBA-1 staining was isolated by splitting fluorescent channels. A max projection was generated from all analyzed images. Next, a median filter was applied to all images. A triangle threshold was applied and this was used to create a mask. The adjustable watershed function with a tolerance of 5 was used to segment individual cells. The analyze particles function was used to measure iMGL area, circularity, cell number, and Feret’s diameter. Only particles with a size greater than 40 were included in the analysis to remove staining from cell debris.

### Meso-Scale Discovery (MSD) V-PLEX Platform to Measure Extracellular Levels of pro and anti-inflammatory Cytokines in iMGL

Pro-and anti-inflammatory cytokines in conditioned media were quantified using the V-PLEX Proinflammatory Panel 1 (Human) kit (#K15049D; Meso Scale Discovery [MSD]) according to the manufacturer’s instructions. Media samples were collected from HBMEC-only, iMGL-only, and iMGL–HBMEC co-cultures described in the “TNFα treatment of 2D co-cultures” section. All media were stored at −80 °C prior to analysis to preserve analyte stability.

For each sample, 50 μL of culture supernatant was loaded into the assay plate. Cytokine levels were detected using the MSD QuickPlex SQ 120 instrument. The assay enabled simultaneous quantification of multiple cytokines, including IL-1β, IL-2, IL-4, IL-6, IL-8, IL-10, IL-12p70, IL-13, IFN-γ, and TNF-α.

### iMGL-Microvessel and Neuroimmune-vascular Model Fabrication

Microvessels were fabricated using a combination of soft lithography and injection molding, following a previously developed protocol^61^. To begin, a 15 mg/mL stock solution of type I collagen (generated from rat tails) was prepared in 0.1% acetic acid. On ice, this collagen solution was neutralized and diluted to a working concentration of 7.5 mg/mL using 1 M sodium hydroxide, 10x M199 medium (Cat# 11825015; Thermo Fisher Scientific), and 1:1 mixture of HBMEC GM and microglia differentiation medium (referred to as 1:1 medium). Following collagen neutralization, iMGLs were lifted from 6 well culture plates using warm Accutase (#07930, STEMCELL Technologies), pelleted, resuspended in 1:1 medium, and mixed into collagen at a concentration of 1M iMGL/mL collagen. A corresponding control collagen gel was made simultaneously, but this control collagen lacked iMGL. Collagen containing iMGLs was used to make iMGL-vessels, whereas control collagen was used to make EC only vessels. With both gels, a PDMS stamp patterned with a 2 channel, hexagon vessel geometry containing a channel diameter of 100 μm was used to mold the collagen into the desired geometry. A separate flat collagen gel (with or without iMGL) was cast and then placed atop the corresponding iMGL or control molded layer after gelation at 37 °C, effectively sealing the microchannel network to form enclosed, perfusable vessels. Thermal incubation at 37 °C facilitated bonding between the two collagen layers.

Once the control or iMGL-microvessels were formed, primary HBMECs were harvested using 0.05% Trypsin-EDTA (Cat# 25300054; Gibco), then resuspended in HBMEC GM at a final concentration of 7-10 × 10⁶ cells/mL. A 10 μL volume of this cell suspension was introduced into the inlet of each vessel using a gel-loading pipette tip. A 10 μL volume of cell suspension was then introduced into the outlet of each vessel. The devices were left under static conditions for 1 hour after cell seeding to allow cells to attach circumferentially along the collagen channel walls. After cell adhesion, gravity-driven flow was initiated. Constructs were cultured under flow conditions for 7 days, with medium exchanged approximately every 12 hours.

### TNFα treatment of 3D engineered microvessels

Control or iMGL-microvessels were treated with TNFα (#H8916, Sigma-Aldrich) after 7 days of culture. The 10 µg/mL TNFα stock was diluted in a 1:1 mixture of HBMEC GM and microglia differentiation medium to a concentration of 0.1 ng/mL. TNFα treatment was performed by adding 100 uL of 0.1 ng/mL TNFα to a well on the top of the acrylic jig that has direct access to the collagen surrounding the vessels. Untreated vessels from the control and microglia-microvessel conditions were also included as additional controls. Vessels with or without TNFα treatment were incubated for 18 hours without disturbance.

### Western Blot

For western blot analysis, cell lysates from iMGL monocultures, iMGL-EC co-cultures, iMGL CM treated cultures, and HBMEC monocultures were collected using 1X RIPA protein lysis buffer (20-188; Millipore) containing 1X protease (#535142-1SET; Millipore) and 1X phosphatase inhibitors (#78427; Thermo Fisher scientific). Protein concentration was quantified using a BSA (#23225; Thermo Fisher Scientific). To run the blot, 20 μg of cell lysates were run on a 4-15% MINI-PROTEAN TGX Precast Protein Gel (#4561084; Bio-Rad) and transferred to a PVDF membrane. 5% BSA in PBS was used for membrane blocking. The membrane was then washed with 1X PBS with 0.05% tween 20 (PBST) and incubated overnight at 4°C with a VE-Cadherin antibody (#ab33168; abcam) diluted 1:1000 in 5% BSA-PBST. The membrane was washed with PBST and incubated with a Goat anti-Rabbit IgG (H+L) Secondary Antibody (#31460; Invitrogen) diluted 1:7500 in 5% milk-PBST for one hour. The membrane was washed with PBST followed by a final PBS wash. The membrane was probed with WesternSure Premium Luminol Enhancer Solution (#826-13460; Li-Cor) and imaged using an Odyssey Clx imaging system (Li-Cor). The membrane was then washed with PBST + 0.2% sodium azide, blocked with 5% milk in PBST + 0.2% sodium azide, and cut at ∼50 kDa. The top portion of the membrane was incubated with ZO-1 antibody (#33-9100; Invitrogen) diluted 1:1000 in 5% BSA-PBST + 0.2% sodium azide overnight at 4°C. The membrane was washed with PBST and incubated with a Goat anti-Mouse IgG (H+L) Secondary Antibody (#31430; Invitrogen) diluted 1:7500 in 5% milk-PBST for one hour. The membrane was washed with PBST followed by a final PBS wash. The membrane was probed with Bio-Rad Clarity Max Western ECL substrate (#1705062; Bio-Rad) and imaged using an Odyssey Clx imaging system (Li-Cor). The bottom portion of the membrane was incubated with GAPDH antibody (#GYX627408; Gene Tex) diluted 1:10,000 in 5% BSA-PBST overnight at 4°C. The membrane was washed with PBST and incubated with a Goat anti-Mouse IgG (H+L) Secondary Antibody (#31430; Invitrogen) diluted 1:7500 in 5% milk-PBST for one hour. The membrane was washed with PBST followed by a final PBS wash. The membrane was probed with Bio-Rad Clarity Western ECL substrate (#1705060; Bio-Rad) and imaged using an Odyssey Clx imaging system (Li-Cor).

### Immunocytochemistry (ICC)

2D cultures of iMGL-HBMEC co-cultures, HBMEC only, and iMGL only were fixed with 4% paraformaldehyde (PFA) for 20 min and washed with PBS three times for 5 mins per wash. Microvessels were fixed by perfusing warm 4% PFA through the microchannels via the inlet and outlet of acrylic jigs using gravity driven flow. This was followed by three 15-20 min washes with PBS also performed via gravity driven flow. Antibody and staining solutions (described below) were perfused through vessels using gravity driven flow in the same way. For both 2D cultures and microvessels, blocking and permeabilization were done simultaneously by treating the cultures with 2% BSA with 0.1% Triton-x for ∼1 hour at room temperature. Next, primary antibodies were resuspended in 2% BSA with 0.1% Triton-X according to the table below (Table 1) and added to fixed samples. Primary antibodies were incubated with samples overnight at 4°C. The following day, 2D cultures were washed with PBS three times for 5 mins, while 3D microvessels were washed three times for 20 mins/wash. Then, secondary antibodies (Table 1) were diluted in 2% BSA (A0281; Sigma) with 0.1% Triton-X 100 (Sigma Aldrich, St Louis, MO) and incubated with the samples overnight at 4°C in the dark. The following day, 2D cultures were washed with PBS three times for 5 mins, while 3D microvessels were washed three times for 20 mins/wash.

**Table 1.**
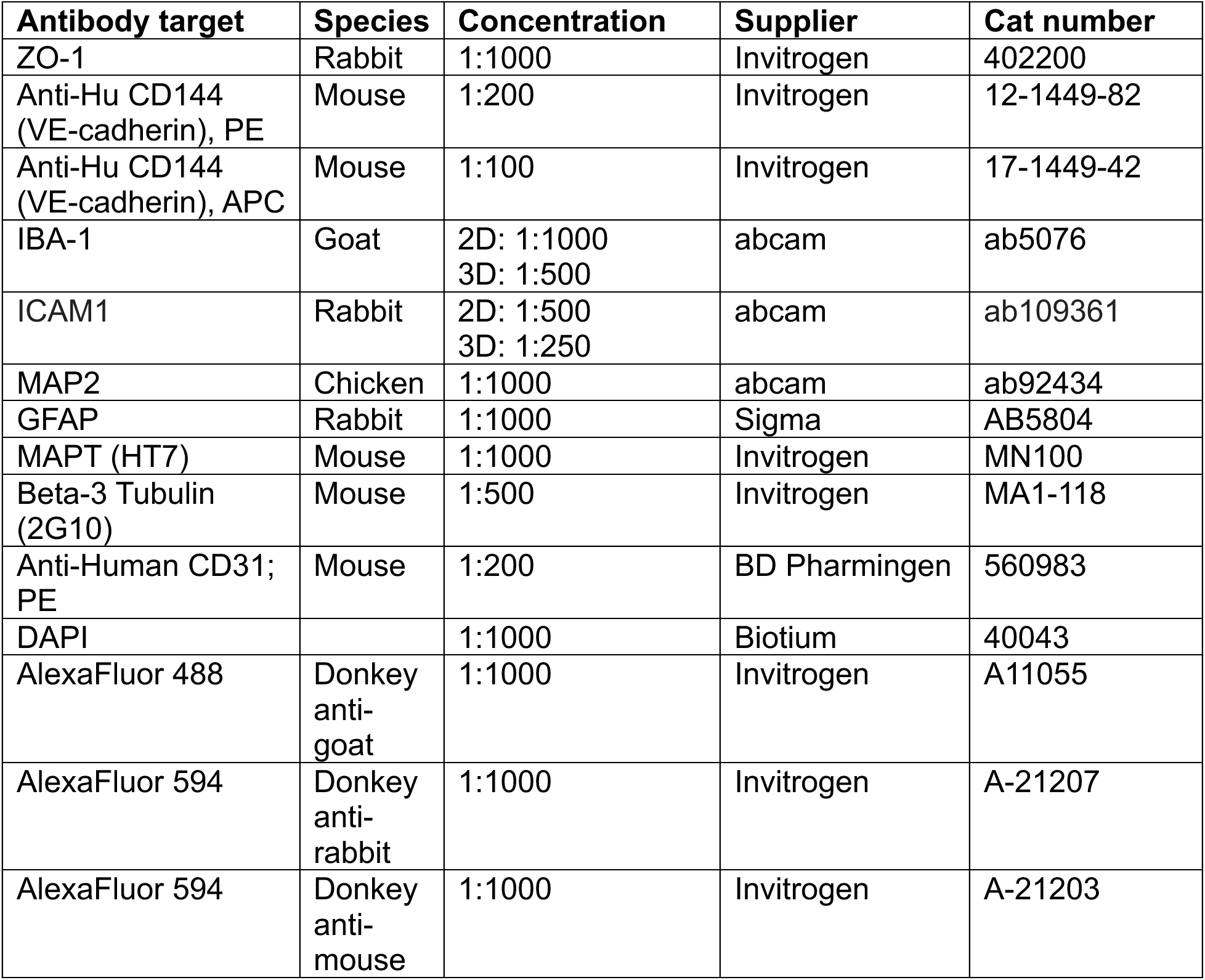
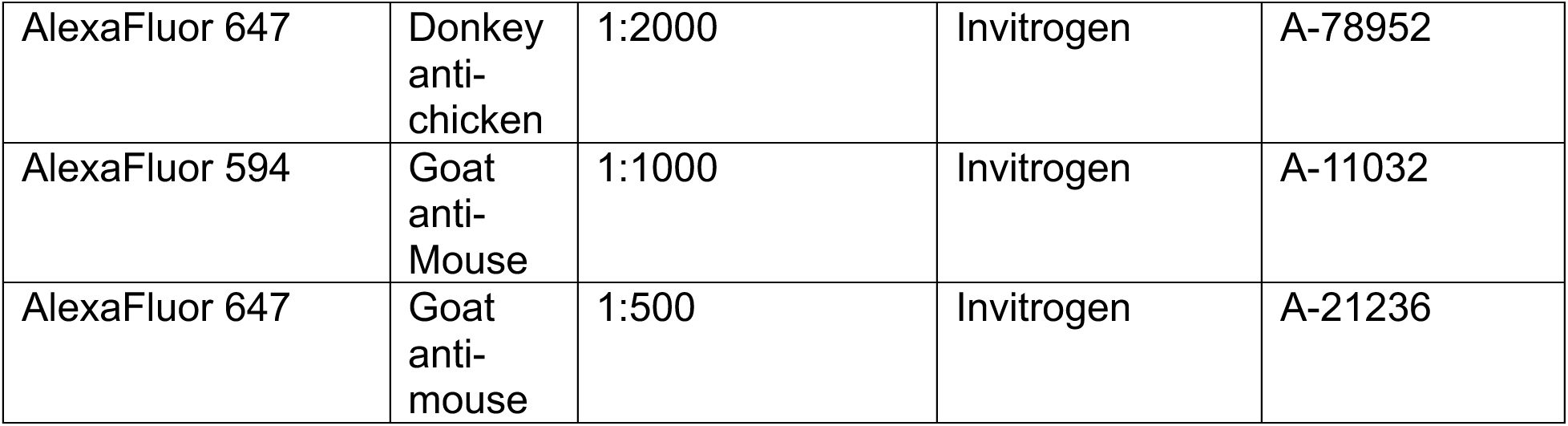
List of antibodies used and their concentrations.

| **Table 1. List of antibodies used and their concentrations.** |  |  |  |  |
| --- | --- | --- | --- | --- |
| **Antibody target** | **Species** | **Concentration** | **Supplier** | **Cat number** |
| ZO-1 | Rabbit | 1:1000 | Invitrogen | 402200 |
| Anti-Hu CD144<br>(VE-cadherin), PE | Mouse | 1:200 | Invitrogen | 12-1449-82 |
| Anti-Hu CD144<br>(VE-cadherin), APC | Mouse | 1:100 | Invitrogen | 17-1449-42 |
| IBA-1 | Goat | 2D: 1:1000<br>3D: 1:500 | abcam | ab5076 |
| ICAM1 | Rabbit | 2D: 1:500<br>3D: 1:250 | abcam | ab109361 |
| MAP2 | Chicken | 1:1000 | abcam | ab92434 |
| GFAP | Rabbit | 1:1000 | Sigma | AB5804 |
| MAPT (HT7) | Mouse | 1:1000 | Invitrogen | MN100 |
| Beta-3 Tubulin<br>(2G10) | Mouse | 1:500 | Invitrogen | MA1-118 |
| Anti-Human CD31;<br>PE | Mouse | 1:200 | BD Pharmingen | 560983 |
| DAPI |  | 1:1000 | Biotium | 40043 |
| AlexaFluor 488 | Donkey<br>anti-<br>goat | 1:1000 | Invitrogen | A11055 |
| AlexaFluor 594 | Donkey<br>anti-rabbit | 1:1000 | Invitrogen | A-21207 |
| AlexaFluor 594 | Donkey<br>anti-mouse | 1:1000 | Invitrogen | A-21203 |
| AlexaFluor 647 | Donkey anti-chicken | 1:2000 | Invitrogen | A-78952 |
| AlexaFluor 594 | Goat anti-Mouse | 1:1000 | Invitrogen | A-11032 |
| AlexaFluor 647 | Goat anti-mouse | 1:500 | Invitrogen | A-21236 |

Cultures of mixed neurons, tricultures of neurons, and microglia only, cultured on coverslips, were fixed with 4% PFA for 15 min and washed with PBS three times for 5 mins per wash. The samples were then stained by first permeabilizing for 20 mins with 0.1% Triton-X 100 (BP151; Fisher). Samples were then washed two times for 5 mins in PBS. The samples were then blocked in 2.5% BSA in PBS for 1 hour at room temperature. Following blocking, the cultures were then stained with primary antibodies at concentrations according to the table below (Table 1) in 2.5% BSA in PBS overnight at 4°C. Next, the samples were washed three times for 5 mins/wash. Secondary antibodies were then diluted according to the table below (Table 1) in 2.5% BSA in PBS overnight at 4°C in the dark. The next day, samples were washed three times in PBS for 5 mins/wash. Lastly, the coverslips were mounted on slides using ProLong Gold Antifade mountant (#P36930; Thermo Fisher Scientific), allowed to dry overnight, sealed with nail polish. Samples were subsequently imaged on a Leica SP8 laser confocal microscope.

### Imaging and Quantification of ICAM1, ZO-1, and VECAD MFI in 2D

Following ICC, 2D cultures of iMGL-HBMEC co-cultures, HBMEC only, and iMGL only were imaged on a Leica SP8 laser confocal microscope using a 25x water immersion objective (HC FLUOTAR L 25x/0,95 W VISIR). For all 2D EC-iMGL co-culture experiments, four 25x images were captured per condition across two technical replicate wells. This was repeated for each biological replicate of this experiment (3 biological replicates total). Z-stack images were captured at a step size of 2.4 μm. To quantify mean fluorescence intensity (MFI) of ICAM1 in TNF⍺ treated cultures and ZO-1, and VECAD MFI in WT co-cultures vs. controls, images were first maximum projected, separated into individual channels to isolate the fluorescence for each protein of interest, and saved as tiff files using ImageJ FIJI Software. Mean grey area was measured by first thresholding the image applying a Huang threshold. Five consistent regions of interest (ROIs) were randomly selected and applied across each image. The mean grey area was then measured and recorded from each ROI. Results of this analysis were plotted using GraphPad Prism software.

### Permeability of engineered microvessels at baseline and after TNFα treatment

The permeability of control and iMGL-containing microvessels to 70 kDa FITC-conjugated dextran (100 μg/mL; [D1823, fisher scientific]) was evaluated both at baseline and following an 18-hour treatment with TNFα. FITC-dextran was prepared in a 1:1 mixture of HBMEC growth medium and microglia differentiation medium and introduced into the microvessel lumen via gravity-driven flow, producing an estimated pressure drop of ∼1 mmHg. Dextran leakage across the endothelial barrier was assessed using time-lapse fluorescence microscopy over a 5-minute interval. Each microvessel device consisted of two parallel channels. Fluorescent images were captured from one channel every second for a 1-minute interval before switching to the second channel for the subsequent minute, alternating between channels for the duration of the 5-minute measurement. Following the TNFα permeability assessment, microvessels were fixed in 4% PFA for 20 minutes at room temperature, then washed three times with PBS in preparation for immunocytochemistry and imaging.

### Quantification of the permeation index

Dextran diffusion across vessel walls was quantified from fluorescent images using Fiji (ImageJ). Measurements of lumen diameter, ROI placement, and fluorescence intensity were extracted and analyzed with custom MATLAB scripts.

A dimensionless metric, the permeation index (Δx), was used to more sensitively capture dextran leakage. In TNFα-injured vessels, high baseline leakage limited the utility of traditional measures. The permeation index was defined as the normalized distance of dextran penetration beyond the luminal boundary. For each vessel, a midpoint ROI was divided into three windows (ci, ci1, ci2; 100 × 1848 pixels), and fluorescence intensity profiles were generated. Within each profile, the x-positions corresponding to one-third and two-thirds of the peak fluorescence intensity were identified, and distances from the lumen center to these points (Δxt, Δxb) were calculated for both vessel walls. The mean of these values across the three windows was then normalized to the lumen diameter, yielding Δx. Lower Δx values correspond to limited dextran spread and an intact barrier, while higher values indicate increased permeability.

### iPSC-derived neuron differentiation

Cortical neurons were generated from the wild-type (WT) control hiPSC line described in the iMGL differentiation section using a previously published dual-SMAD inhibition protocol^62, 64, 65^. For the experiments described here, previously differentiated and cryopreserved neural precursor cells (NPCs) were thawed and used as the starting population.

NPCs were cultured in 6-well plates coated with Geltrex™ LDEV-Free Reduced Growth Factor Basement Membrane Matrix (#A1413202; Gibco) and maintained in Basal Neural Maintenance Medium (BNMM) supplemented with fibroblast growth factor (FGF). BNMM consisted of a 1:1 mixture of DMEM/F12 with Glutamine (#11320033; Gibco) and Neurobasal medium (#21103049; Gibco), supplemented with 0.5% N2 (#17502048; Gibco), 1% B27 (#17504044; Gibco), 0.5% GlutaMAX (#35050061; Thermo Fisher Scientific), 0.5% insulin-transferrin-selenium (#41400045; Thermo Fisher Scientific), 0.5% non-essential amino acids (NEAA; #11140050; Thermo Fisher Scientific), 0.2% β-mercaptoethanol (#21985023; Life Technologies), and 20 ng/mL FGF (R&D Systems). Cells were fed every other day and grown to ∼90% confluency.

Confluent NPCs were dissociated using Accutase (#07930; STEMCELL Technologies) and replated onto Geltrex-coated 10 cm dishes for neuronal differentiation. Upon reaching ∼90% confluency, cells were transitioned to neural differentiation medium, consisting of BNMM supplemented with 0.2 µg/mL brain-derived neurotrophic factor (BDNF; #450-02; PeproTech), 0.2 µg/mL glial cell-derived neurotrophic factor (GDNF; #450-10; PeproTech), and 0.5 mM dibutyryl-cAMP (dbcAMP; #D0260; Sigma-Aldrich). Cells were maintained in this medium for 21 days with biweekly media changes.

Differentiated hiPSC-derived neurons were used in downstream experiments and maintained at 37 °C in a humidified incubator with 5% CO₂.

### Neuron-microglia 2D co-cultures

Following iPSC-derived neuron differentiation, neurons were lifted with warm Accutase (#07930; STEMCELL Technologies) for 30 mins. Neurons were dissociated into a single-cell suspension using trituration with a p1000 pipet. They were then pelleted, counted for live cells using Trypan Blue (#15250061; Gibco), and resuspended at a concentration of 2×10^6^ cells/mL in neural differentiation medium + 10 µM ROCK inhibitor Y-27632 (#A3008; ApexBio). Cells were plated at a density of 200,000 live cells/well in a 24-well plate containing Geltrex™ LDEV-Free Reduced Growth Factor Basement Membrane Matrix (#A1413202; Gibco) coated coverslips to allow for later staining and imaging. The next day, neural differentiation medium + ROCK inhibitor was replaced with neural differentiation medium and neurons were allowed to recover for ∼24 hours.

iMGL were thawed, counted, and plated in 1:60 growth factor-reduced Matrigel (#356231; Corning) coated 6-well plates at a density of 2×10^6^ cells/well in microglia differentiation medium. iMGL were allowed to recover from thawing for 3-5 days prior to use in co-cultures.

Once iMGL and neurons had recovered, iMGL were collected and lifted using Accutase for 5 mins. They were then pelleted and resuspended in 1:1 neuron:microglia medium (BNMM with a base of DMEM-F12 with no phenol red (#11039021; Gibco) (rather than 1:1 mixture of DMEM/F12 with Glutamine (#11320033; Gibco) and Neurobasal medium (#21103049; Gibco)) supplemented with BDNF, GDNF, and dcAMP at a 1:1 ratio with microglia differentiation medium). Neuron media on the neurons was also replaced with 1:1 neuron:microglia medium. iMGL were then added to the wells containing microglia at a density of 200k cells/well. Controls for these cultures include neuron only and microglia only controls at the same densities in the same 1:1 culture medium. Neuron-iMGL co-cultures were fed through media addition on alternate days for 7 days, after which co-cultures and their corresponding controls were fixed with 4% PFA for 15 mins and then washed with PBS three times for 5 mins each for subsequent ICC and imaging.

### B3tub+ and MAP2+ branch length quantification

Following ICC, 2D cultures of mixed culture neurons – including neurons and astrocytes, tricultures, and iMGL only were imaged on a Leica SP8 laser confocal microscope using a 20x objective. For all 2D co-culture experiments, four 20x images were captured per condition across two technical replicate coverslips. This was repeated for each biological replicate of this experiment (3 biological replicates total). Z-stack images were captured. To quantify MAP2+ branch length in the 2D co-cultures we used custom ImageJ and python scripts. The process was as follows: the fluorescent channels were split so that MAP2 signal was isolated, a maximum intensity projection was made, each image was skeletonized, segmented, and the segments were labelled. Next, the total number of segments was calculated, the total length of segments was calculated, and then the total length was divided by the number of segments to determine the average segment length across the image. This was reported as the branch length of the neurites.

In 3D NVU and neuron vessels, 10x images were captured on a Leica SP8 laser confocal microscope. Three images per vessel were taken across 4-8 biological replicates. Z-stack images were captured. B3TUB+ branch length was calculated using the same method described above. However, before skeletonization, outliers were removed from the image to limit the influence of cell debris on our calculations.

### Multielectrode array (MEA) on tricultures

Mixed neuronal cultures generated as described above were replated onto microelectrode array (MEA) plates coated with Matrigel. Both 6-well and 48-well plate formats were used in these studies (#M384-tMEA-6W, #M768-tMEA-48W; Axion Biosystems). The Matrigel coating was applied selectively over the electrode region, and residual matrix solution was removed immediately before seeding. Cells were deposited at a density of 10,000 cells/µl, corresponding to ∼300,000 neurons per well in the 6-well format and ∼100,000 neurons per well in the 48-well format. Following plating, cultures were maintained for the 48 hours in BNMM medium supplemented with BDNF, GDNF, and db-cAMP (see above). After 48 hours, iMGL that had had been cultured for 3-5 days post differentiation were lifted using accutase and added to the MEA culture wells at a 1:1 ratio with the neurons and astrocytes. At this point the medium on the cultures was transitioned to BrainPhys without phenol red (#05791; Stem Cell Technologies). The BrainPhys was supplemented with 1% B27, 0.5% N2, 0.1 μg/ml BDNF, 0.1 μg/ml GDNF, 250 μM db-cAMP, mCSF, TGF-β, IL-34, and 1 μg/ml laminin to support neurons and iMGL and promote electrical activity in the cultures. Media were partially exchanged twice weekly, with two-thirds of the total well volume replaced at each change.

### MEA recordings and analysis

Extracellular activity of hiPSC-derived neurons cultured on MEA plates was monitored twice weekly using the Maestro Pro system (Axion Biosystems). Recordings were performed prior to media changes to minimize mechanical disturbance. Plates were equilibrated in the MEA reader for roughly 10 minutes before starting recordings, with temperature and CO₂ maintained at 37°C and 5%, respectively. Each recording lasted approximately five minutes. Neuronal spikes were identified using the Axion Axis Navigator software. Electrodes registering fewer than five spikes per minute were excluded from subsequent analyses. For metric calculations, a three-minute segment of each recording was used. Mean firing rate and network burst frequency were quantified using Axion’s Neural Metric Tool software. Network bursts were defined using an adaptive threshold, with a minimum of 35% of electrodes required to participate for a burst to be counted. Raster plots and spike histograms were generated using custom MATLAB scripts.

## Statistical Analysis

Data were first assessed for normality using the Shapiro–Wilk test. For datasets meeting the assumptions of normality, statistical comparisons were performed using two-tailed unpaired t-tests, one-way ANOVA, or two-way ANOVA as appropriate. Differences were considered statistically significant at p < 0.05. All analyses were conducted using GraphPad Prism software. Information on statistical details of individual experiments can be found in the respective figure legends.

## List of Abbreviations

AD: Alzheimer’s disease
2D: Two-dimensional
3D: Three-dimensional
NVU: Neurovascular unit
iNs: Human induced pluripotent stem cell derived-neurons
iAs: Human induced pluripotent stem cell derived-astrocytes
iMGLs: Human induced pluripotent stem cell derived-microglia-like cells
HBMECs: Human brain microvascular endothelial cells
EC: Endothelial cell
TNFα: Tumor necrosis factor α
BBB: Blood brain barrier
CNS: Central nervous system
IL: Interleukin
GWAS: Genome wide association studies
TEER: Transendothelial electrical resistance
iPSC: human induced pluripotent stem cell
LPS: lipopolysaccharide
CSF-1R: Colony-Stimulating Factor 1 Receptor
ZO-1: zonula occludens-1
B3TUB: βIII-tubulin
ICC: Immunocytochemistry
MAP2: microtubule associated protein 2
MAPT: microtubule associated protein tau
GFAP: glial fibrillary acidic protein
IBA-1: ionized calcium-binding adaptor molecule 1
MEA: multielectrode array
MFI: mean fluorescence intensity
VECAD: VE-Cadherin
CM: conditioned media CSF: cerebrospinal fluid
ICAM1: intercellular adhesion molecule 1
MSD: Mesoscale Discovery

## References

1. Sweeney MD, Kisler K, Montagne A, Toga AW, Zlokovic B V. (2018) The role of brain vasculature in neurodegenerative disorders. Nat Neurosci 21:1318

2. Soto-Rojas LO, Pacheco-Herrero M, Martínez-Gómez PA, Campa-Córdoba BB, Apátiga-Pérez R, Villegas-Rojas MM, Harrington CR, de la Cruz F, Garcés-Ramírez L, Luna-Muñoz J (2021) The Neurovascular Unit Dysfunction in Alzheimer’s Disease. Int J Mol Sci 22:2022

3. Hawkins BT, Davis TP (2005) The Blood-Brain Barrier/Neurovascular Unit in Health and Disease. Pharmacol Rev 57:173–185

4. Daneman R (2012) The blood-brain barrier in health and disease. Ann Neurol 72:648–672

5. Zlokovic B V. (2008) The blood-brain barrier in health and chronic neurodegenerative disorders. Neuron 57:178–201

6. Lochhead JJ, Yang J, Ronaldson PT, Davis TP (2020) Structure, Function, and Regulation of the Blood-Brain Barrier Tight Junction in Central Nervous System Disorders. Front Physiol 11:558491

7. Zlokovic B V. (2011) Neurovascular pathways to neurodegeneration in Alzheimer’s disease and other disorders. Nat Rev Neurosci 12:723

8. Iadecola C (2013) The Pathobiology of Vascular Dementia. Neuron 80:844–866

9. Snyder HM, Corriveau RA, Craft S, et al (2015) Vascular contributions to cognitive impairment and dementia including Alzheimer’s disease. Alzheimer’s & Dementia 11:710–717

10. Iadecola C, Davisson RL (2008) Hypertension and Cerebrovascular Dysfunction. Cell Metab 7:476–484

11. Toledo JB, Arnold SE, Raible K, Brettschneider J, Xie SX, Grossman M, Monsell SE, Kukull WA, Trojanowski JQ (2013) Contribution of cerebrovascular disease in autopsy confirmed neurodegenerative disease cases in the National Alzheimer’s Coordinating Centre. Brain 136:2697

12. Montagne A, Barnes SR, Sweeney MD, et al (2015) Blood-Brain barrier breakdown in the aging human hippocampus. Neuron 85:296–302

13. Nation DA, Sweeney MD, Montagne A, et al (2019) Blood-brain barrier breakdown is an early biomarker of human cognitive dysfunction. Nat Med 25:270–276

14. Gorelick PB, Scuteri A, Black SE, et al (2011) Vascular contributions to cognitive impairment and dementia: A statement for healthcare professionals from the American Heart Association/American Stroke Association. Stroke 42:2672–2713

15. Zipser BD, Johanson CE, Gonzalez L, Berzin TM, Tavares R, Hulette CM, Vitek MP, Hovanesian V, Stopa EG (2007) Microvascular injury and blood–brain barrier leakage in Alzheimer’s disease. Neurobiol Aging 28:977–986

16. Yamazaki Y, Kanekiyo T (2017) Blood-Brain Barrier Dysfunction and the Pathogenesis of Alzheimer’s Disease. International Journal of Molecular Sciences 2017, Vol 18, Page 1965 18:1965

17. Yu X, Ji C, Shao A (2020) Neurovascular Unit Dysfunction and Neurodegenerative Disorders. Front Neurosci 14:534494

18. Montagne A, Nation DA, Sagare AP, et al (2020) APOE4 leads to blood-brain barrier dysfunction predicting cognitive decline. Nature 581:71–76

19. Nelson AR (2022) Peripheral Pathways to Neurovascular Unit Dysfunction, Cognitive Impairment, and Alzheimer’s Disease. Front Aging Neurosci 14:858429

20. Govindpani K, McNamara LG, Smith NR, Vinnakota C, Waldvogel HJ, Faull RLM, Kwakowsky A (2019) Vascular Dysfunction in Alzheimer’s Disease: A Prelude to the Pathological Process or a Consequence of It? Journal of Clinical Medicine 2019, Vol 8, Page 651 8:651

21. Wang D, Chen F, Han Z, Yin Z, Ge X, Lei P (2021) Relationship Between Amyloid-β Deposition and Blood–Brain Barrier Dysfunction in Alzheimer’s Disease. Front Cell Neurosci 15:695479

22. Argaw AT, Asp L, Zhang J, et al (2012) Astrocyte-derived VEGF-A drives blood-brain barrier disruption in CNS inflammatory disease. J Clin Invest 122:2454–2468

23. McGeer PL, Itagaki S, Tago H, McGeer EG (1987) Reactive microglia in patients with senile dementia of the Alzheimer type are positive for the histocompatibility glycoprotein HLA-DR. Neurosci Lett 79:195–200

24. Davies P, Maloney AJF (1976) Selective loss of central cholinergic neurons in Alzheimer’s disease. Lancet 2:1403

25. Matthews KL, Chen CPLH, Esiri MM, Keene J, Minger SL, Francis PT (2002) Noradrenergic changes, aggressive behavior, and cognition in patients with dementia. Biol Psychiatry 51:407–416

26. Šimić G, Babić Leko M, Wray S, et al (2017) Monoaminergic neuropathology in Alzheimer’s disease. Prog Neurobiol 151:101–138

27. Li Q, Barres BA (2017) Microglia and macrophages in brain homeostasis and disease. Nature Reviews Immunology 2017 18:4 18:225–242

28. Tremblay MÈ, Stevens B, Sierra A, Wake H, Bessis A, Nimmerjahn A (2011) The Role of Microglia in the Healthy Brain. Journal of Neuroscience 31:16064–16069

29. Colonna M, Butovsky O (2017) Microglia Function in the Central Nervous System During Health and Neurodegeneration. Annu Rev Immunol 35:441

30. Bachiller S, Jiménez-Ferrer I, Paulus A, Yang Y, Swanberg M, Deierborg T, Boza-Serrano A (2018) Microglia in neurological diseases: A road map to brain-disease dependent-inflammatory response. Front Cell Neurosci 12:403344

31. McQuade A, Blurton-Jones M (2019) Microglia in Alzheimer’s Disease: Exploring How Genetics and Phenotype Influence Risk. J Mol Biol 431:1805–1817

32. Bertram L, Lange C, Mullin K, et al (2008) Genome-wide Association Analysis Reveals Putative Alzheimer’s Disease Susceptibility Loci in Addition to APOE. Am J Hum Genet 83:623

33. Hollingworth P, Harold D, Sims R, et al (2011) Common variants in ABCA7, MS4A6A/MS4A4E, EPHA1, CD33 and CD2AP are associated with Alzheimer’s disease. Nat Genet 43:429

34. Naj AC, Jun G, Beecham GW, et al (2011) Common variants in MS4A4/MS4A6E, CD2uAP, CD33, and EPHA1 are associated with late-onset Alzheimer’s disease. Nat Genet 43:436

35. Jonsson T, Stefansson H, Steinberg S, et al (2012) Variant of TREM2 Associated with the Risk of Alzheimer’s Disease. N Engl J Med 368:107

36. Lambert JC, Zelenika D, Hiltunen M, et al (2011) Evidence of the association of BIN1 and PICALM with the AD risk in contrasting European populations. Neurobiol Aging 32:756.e11–756.e15

37. Seshadri S, Fitzpatrick AL, Ikram MA, et al (2010) Genome-wide Analysis of Genetic Loci Associated with Alzheimer’s Disease. JAMA: the journal of the American Medical Association 303:1832

38. Griciuc A, Tanzi RE (2021) The role of innate immune genes in Alzheimer’s disease. Curr Opin Neurol 34:228

39. Friedman BA, Srinivasan K, Ayalon G, et al (2018) Diverse Brain Myeloid Expression Profiles Reveal Distinct Microglial Activation States and Aspects of Alzheimer’s Disease Not Evident in Mouse Models. Cell Rep 22:832–847

40. Ueda Y, Gullipalli D, Song WC (2016) Modeling complement-driven diseases in transgenic mice: Values and limitations. Immunobiology 221:1080–1090

41. Booth R, Kim H (2012) Characterization of a microfluidic in vitro model of the blood-brain barrier (μBBB). Lab Chip 12:1784–1792

42. Stone NL, England TJ, O’Sullivan SE (2019) A novel transwell blood brain barrier model using primary human cells. Front Cell Neurosci 13:455689

43. Williams-Medina A, Deblock M, Janigro D (2021) In vitro Models of the Blood–Brain Barrier: Tools in Translational Medicine. Front Med Technol 2:623950

44. Brown JA, Pensabene V, Markov DA, et al (2015) Recreating blood-brain barrier physiology and structure on chip: A novel neurovascular microfluidic bioreactor. Biomicrofluidics 9:054124

45. Pediaditakis I, Kodella KR, Manatakis D V., et al (2021) Modeling alpha-synuclein pathology in a human brain-chip to assess blood-brain barrier disruption. Nature Communications 2021 12:1 12:1–17

46. Ko E, Kamm RD (2022) Neurovascular models for organ-on-a-chips. In vitro models 2022 1:2 1:125–127

47. Adriani G, Ma D, Pavesi A, Kamm RD, Goh ELK (2017) A 3D neurovascular microfluidic model consisting of neurons, astrocytes and cerebral endothelial cells as a blood–brain barrier. Lab Chip 17:448–459

48. Shin Y, Choi SH, Kim E, Bylykbashi E, Kim JA, Chung S, Kim DY, Kamm RD, Tanzi RE (2019) Blood–Brain Barrier Dysfunction in a 3D In Vitro Model of Alzheimer’s Disease. Advanced Science 6:1900962

49. Kim J, Lee KT, Lee JS, et al (2021) Fungal brain infection modelled in a human-neurovascular-unit-on-a-chip with a functional blood–brain barrier. Nature Biomedical Engineering 2021 5:8 5:830–846

50. Lyu Z, Park J, Kim KM, Jin HJ, Wu H, Rajadas J, Kim DH, Steinberg GK, Lee W (2021) A neurovascular-unit-on-a-chip for the evaluation of the restorative potential of stem cell therapies for ischaemic stroke. Nature Biomedical Engineering 2021 5:8 5:847–863

51. Pediaditakis I, Kodella KR, Manatakis D V., et al (2022) A microengineered Brain-Chip to model neuroinflammation in humans. iScience. 10.1016/J.ISCI.2022.104813

52. Bisht K, Okojie KA, Sharma K, et al (2021) Capillary-associated microglia regulate vascular structure and function through PANX1-P2RY12 coupling in mice. Nature Communications 2021 12:1 12:1–13

53. Mondo E, Becker SC, Kautzman AG, Schifferer M, Baer CE, Chen J, Huang EJ, Simons M, Schafer DP (2020) A Developmental Analysis of Juxtavascular Microglia Dynamics and Interactions with the Vasculature. Journal of Neuroscience 40:6503–6521

54. Haruwaka K, Ikegami A, Tachibana Y, et al (2019) Dual microglia effects on blood brain barrier permeability induced by systemic inflammation. Nature Communications 2019 10:1 10:1–17

55. Sumi N, Nishioku T, Takata F, Matsumoto J, Watanabe T, Shuto H, Yamauchi A, Dohgu S, Kataoka Y (2009) Lipopolysaccharide-Activated Microglia Induce Dysfunction of the Blood–Brain Barrier in Rat Microvascular Endothelial Cells Co-Cultured with Microglia. Cell Mol Neurobiol 30:247

56. Bull D, Schweitzer C, Bichsel C, Britschgi M, Gutbier S (2022) Generation of an hiPSC-Derived Co-Culture System to Assess the Effects of Neuroinflammation on Blood–Brain Barrier Integrity. Cells 11:419

57. Profaci CP, Harvey SS, Bajc K, et al (2024) Microglia are not necessary for maintenance of blood-brain barrier properties in health, but PLX5622 alters brain endothelial cholesterol metabolism. Neuron 112:2910–2921.e7

58. Elmore MRP, Najafi AR, Koike MA, et al (2014) Colony-Stimulating Factor 1 Receptor Signaling Is Necessary for Microglia Viability, Unmasking a Microglia Progenitor Cell in the Adult Brain. Neuron 82:380–397

59. Delaney C, Farrell M, Doherty CP, et al (2021) Attenuated CSF-1R signalling drives cerebrovascular pathology. EMBO Mol Med. https://doi.org/10.15252/EMMM.202012889/SUPPL_FILE/EMMM202012889-SUP-0002-EVFIGS.PDF

60. Mehrabadi AR, Korolainen MA, Odero G, Miller DW, Kauppinen TM (2017) Poly(ADP-ribose) polymerase-1 regulates microglia mediated decrease of endothelial tight junction integrity. Neurochem Int 108:266–271

61. Zheng Y, Chen J, Craven M, et al (2012) In vitro microvessels for the study of angiogenesis and thrombosis. Proc Natl Acad Sci U S A 109:9342–9347

62. Shin YJ, Evitts KM, Jin S, Howard C, Sharp-Milgrom M, Schwarze-Taufiq T, Kinoshita C, Young JE, Zheng Y (2023) Amyloid beta peptides (Aβ) from Alzheimer’s disease neuronal secretome induce endothelial activation in a human cerebral microvessel model. Neurobiol Dis 181:106125

63. McQuade A, Coburn M, Tu CH, Hasselmann J, Davtyan H, Blurton-Jones M (2018) Development and validation of a simplified method to generate human microglia from pluripotent stem cells. Mol Neurodegener. 10.1186/S13024-018-0297-X

64. Knupp A, Mishra S, Martinez R, Braggin JE, Szabo M, Kinoshita C, Hailey DW, Small SA, Jayadev S, Young JE (2020) Depletion of the AD Risk Gene SORL1 Selectively Impairs Neuronal Endosomal Traffic Independent of Amyloidogenic APP Processing. Cell Rep 31:107719

65. Rose SE, Frankowski H, Knupp A, et al (2018) Leptomeninges-Derived Induced Pluripotent Stem Cells and Directly Converted Neurons From Autopsy Cases With Varying Neuropathologic Backgrounds. J Neuropathol Exp Neurol 77:353

66. Kok LML, Helwegen K, Coveña NF, Heine VM (2025) Human pluripotent stem cell-derived microglia shape neuronal morphology and enhance network activity in vitro. J Neurosci Methods 415:110354

67. Lou N, Takano T, Pei Y, Xavier AL, Goldman SA, Nedergaard M (2016) Purinergic receptor P2RY12-dependent microglial closure of the injured blood-brain barrier. Proc Natl Acad Sci U S A 113:1074–1079

68. Vidal-Itriago A, Radford RAW, Aramideh JA, Maurel C, Scherer NM, Don EK, Lee A, Chung RS, Graeber MB, Morsch M (2022) Microglia morphophysiological diversity and its implications for the CNS. Front Immunol 13:997786

69. Caldeira C, Cunha C, Vaz AR, Falcão AS, Barateiro A, Seixas E, Fernandes A, Brites D (2017) Key aging-associated alterations in primary microglia response to beta-amyloid stimulation. Front Aging Neurosci 9:277

70. Fillit H, Ding W, Buee L, Kalman J, Altstiel L, Lawlor B, Wolf-Klein G (1991) Elevated circulating tumor necrosis factor levels in Alzheimer’s disease. Neurosci Lett 129:318–320

71. Swardfager W, Lanctt K, Rothenburg L, Wong A, Cappell J, Herrmann N (2010) A meta-analysis of cytokines in Alzheimer’s disease. Biol Psychiatry 68:930–941

72. Decourt B, Lahiri DK, Sabbagh MN (2017) Targeting Tumor Necrosis Factor Alpha for Alzheimer’s Disease. Curr Alzheimer Res 14:412–425

73. Tarkowski E, Andreasen N, Tarkowski A, Blennow K (2003) Intrathecal inflammation precedes development of Alzheimer’s disease. J Neurol Neurosurg Psychiatry 74:1200–1205

74. Brosseron F, Krauthausen M, Kummer M, Heneka MT (2014) Body fluid cytokine levels in mild cognitive impairment and Alzheimer’s disease: a comparative overview. Mol Neurobiol 50:534–544

75. Holmes C, Cunningham C, Zotova E, Woolford J, Dean C, Kerr S, Culliford D, Perry VH (2009) Systemic inflammation and disease progression in Alzheimer disease. Neurology 73:768–774

76. Dyne E, Cawood M, Suzelis M, Russell R, Kim M-H (2021) Ultrastructural Analysis of the Morphological Phenotypes of Microglia Associated with Neuro-inflammatory Cues. J Comp Neurol 530:1263

77. Versele R, Sevin E, Gosselet F, Fenart L, Candela P (2022) TNF-α and IL-1β Modulate Blood-Brain Barrier Permeability and Decrease Amyloid-β Peptide Efflux in a Human Blood-Brain Barrier Model. International Journal of Molecular Sciences 2022, Vol 23, Page 10235 23:10235

78. Cheng Y, Desse S, Martinez A, Worthen RJ, Jope RS, Beurel E (2018) TNFα disrupts blood brain barrier integrity to maintain prolonged depressive-like behavior in mice. Brain Behav Immun 69:556–567

79. Zhang M, Wang P, Wu Y, et al (2025) A microengineered 3D human neurovascular unit model to probe the neuropathogenesis of herpes simplex encephalitis. Nature Communications 2025 16:1 16:1–16

80. Lloyd AF, Martinez-Muriana A, Davis E, et al (2024) Deep proteomic analysis of microglia reveals fundamental biological differences between model systems. Cell Rep 43:114908

81. Chen WT, Lu A, Craessaerts K, et al (2020) Spatial Transcriptomics and In Situ Sequencing to Study Alzheimer’s Disease. Cell 182:976–991.e19

82. Zhou Y, Song WM, Andhey PS, et al (2020) Human and mouse single-nucleus transcriptomics reveal TREM2-dependent and TREM2-independent cellular responses in Alzheimer’s disease. Nature Medicine 2020 26:1 26:131–142

83. Berry BJ, Smith AST, Young JE, Mack DL (2019) Advances and Current Challenges Associated with the Use of Human Induced Pluripotent Stem Cells in Modeling Neurodegenerative Disease. Cells Tissues Organs 205:331–349

84. Colonna M, Butovsky O (2017) Microglia Function in the Central Nervous System During Health and Neurodegeneration. Annu Rev Immunol 35:441

85. Guzmán-Ruíz MA, Guerrero Vargas NN, Ramírez-Carreto RJ, González-Orozco JC, Torres-Hernández BA, Valle-Rodríguez M, Guevara-Guzmán R, Chavarría A (2024) Microglia in physiological conditions and the importance of understanding their homeostatic functions in the arcuate nucleus. Front Immunol 15:1392077

86. Bozkurt A, Brook GA, Moellers S, et al (2007) In vitro assessment of axonal growth using dorsal root ganglia explants in a novel three-dimensional collagen matrix. Tissue Eng 13:2971–2979

87. Deister C, Aljabari S, Schmidt CE (2007) Effects of collagen 1, fibronectin, laminin and hyaluronic acid concentration in multi-component gels on neurite extension. J Biomater Sci Polym Ed 18:983–997

88. Rusin D, Vahl Becirovic L, Lyszczarz G, Krueger M, Benmamar-Badel A, Vad Mathiesen C, Sigurðardóttir Schiöth E, Lykke Lambertsen K, Wlodarczyk A (2024) Microglia-Derived Insulin-like Growth Factor 1 Is Critical for Neurodevelopment. Cells 13:184

89. Pons V, Rivest S (2020) Beneficial Roles of Microglia and Growth Factors in MS, a Brief Review. Front Cell Neurosci 14:284

90. Bake S, Okoreeh AK, Alaniz RC, Sohrabji F (2015) Insulin-Like Growth Factor (IGF)-I Modulates Endothelial Blood-Brain Barrier Function in Ischemic Middle-Aged Female Rats. Endocrinology 157:61

91. Higashi Y, Sukhanov S, Shai SY, et al (2020) Endothelial deficiency of insulin-like growth factor-1 receptor reduces endothelial barrier function and promotes atherosclerosis in Apoe-deficient mice. Am J Physiol Heart Circ Physiol 319:H730

92. Heithoff BP, George KK, Phares AN, Zuidhoek IA, Munoz-Ballester C, Robel S (2020) Astrocytes are necessary for blood-brain barrier maintenance in the adult mouse brain. Glia 69:436

93. Serafini S, Ferretti G, Monterosso P, Angiolillo A, Di Costanzo A, Matrone C (2024) TNF-α Levels Are Increased in Patients with Subjective Cognitive Impairment and Are Negatively Correlated with β Amyloid-42. Antioxidants 13:216

94. Xue M, Qiqige C, Zhang Q, Zhao H, Su L, Sun P, Zhao P (2018) Effects of Tumor Necrosis Factor α (TNF-α) and Interleukina 10 (IL-10) on Intercellular Cell Adhesion Molecule-1 (ICAM-1) and Cluster of Differentiation 31 (CD31) in Human Coronary Artery Endothelial Cells. Med Sci Monit 24:4433

95. Krakauer T (1995) IL-10 inhibits the adhesion of leukocytic cells to IL-1-activated human endothelial cells. Immunol Lett 45:61–65

96. Lin R, Chen F, Wen S, Teng T, Pan Y, Huang H (2018) Interleukin-10 attenuates impairment of the blood-brain barrier in a severe acute pancreatitis rat model. J Inflamm (Lond) 15:4

97. Worzfeld T, Schwaninger M (2015) Apicobasal polarity of brain endothelial cells. Journal of Cerebral Blood Flow & Metabolism 36:340

98. Lam D, Enright HA, Cadena J, Peters SKG, Sales AP, Osburn JJ, Soscia DA, Kulp KS, Wheeler EK, Fischer NO (2019) Tissue-specific extracellular matrix accelerates the formation of neural networks and communities in a neuron-glia co-culture on a multi-electrode array. Scientific Reports 2019 9:1 9:1–15

99. Zhang W, Xiao D, Mao Q, Xia H (2023) Role of neuroinflammation in neurodegeneration development. Signal Transduction and Targeted Therapy 2023 8:1 8:1–32

100. Mishra S, Morshed N, Sidhu SB, Kinoshita C, Stevens B, Jayadev S, Young JE (2025) The Alzheimer’s Disease Gene SORL1 Regulates Lysosome Function in Human Microglia. Glia 73:1329–1348

101. Levy S, Sutton G, Ng PC, et al (2007) The Diploid Genome Sequence of an Individual Human. PLoS Biol 5:e254

