## Supplemental Data for "Evaluating Microglial Contributions to the Neurovascular Unit in Health and Neurodegeneration Using Human *In Vitro* Models"

**Supplemental Figure 1.**

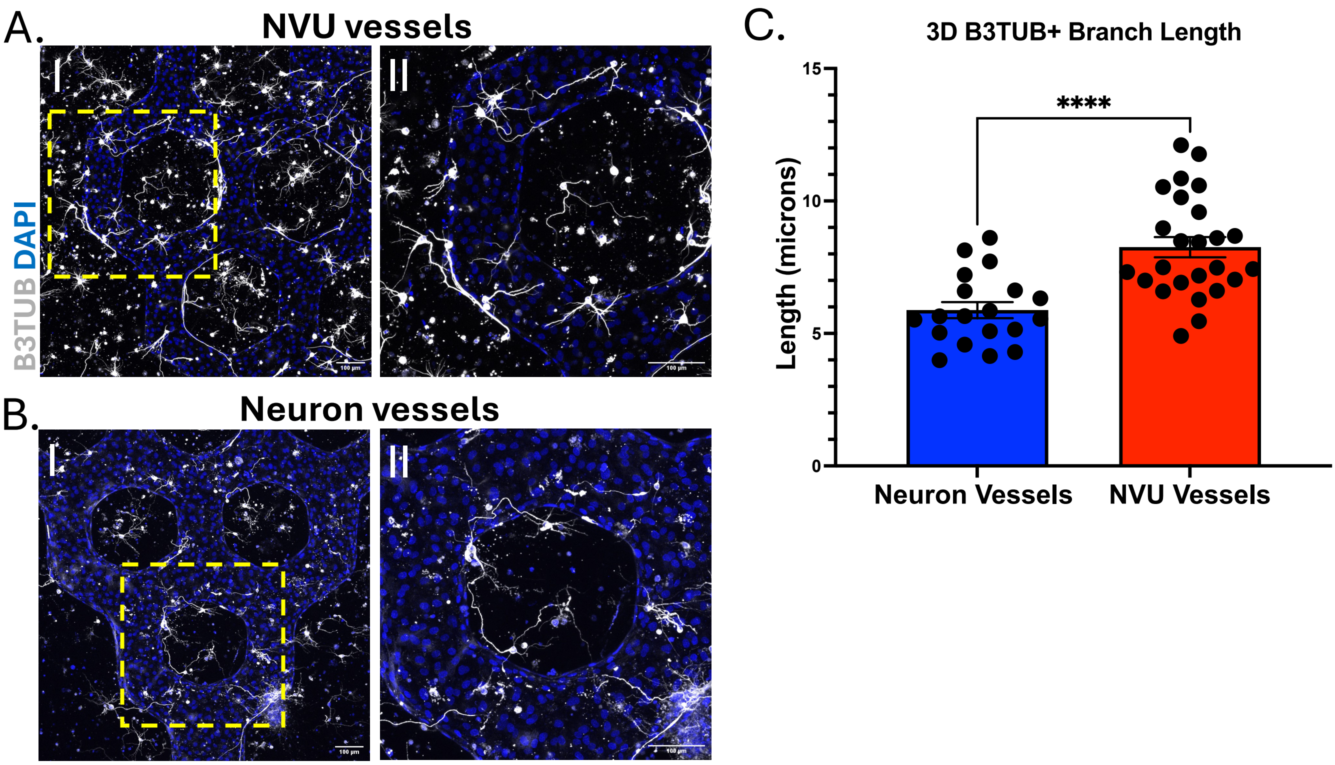


**Supplemental Figure 1. The addition of iMGL in 3D NVU vessels improves neuronal morphology.** (A) Representative I) 10x and II) 20x images of NVU vessels containing neurons, astrocytes, and iMGL. Scale bar = 100 μm. (B) Representative I) 10x and II) 20x images of neuron vessels containing neurons and astrocytes. Neurons in NVU vessels show stronger B3TUB staining (white) compared to neuron vessels. Scale bar = 100 μm. (E) Quantification of B3TUB+ neuronal branch length in NVU vessels vs. neuron vessels. NVU vessels have significantly longer branches compared to neuron vessels. Error bars, mean ± SEM. *p < 0.05, **p < 0.002, ***p < 0.0002, ****p<0.0001 by unpaired t-test.

#### Supplemental Figure 2.

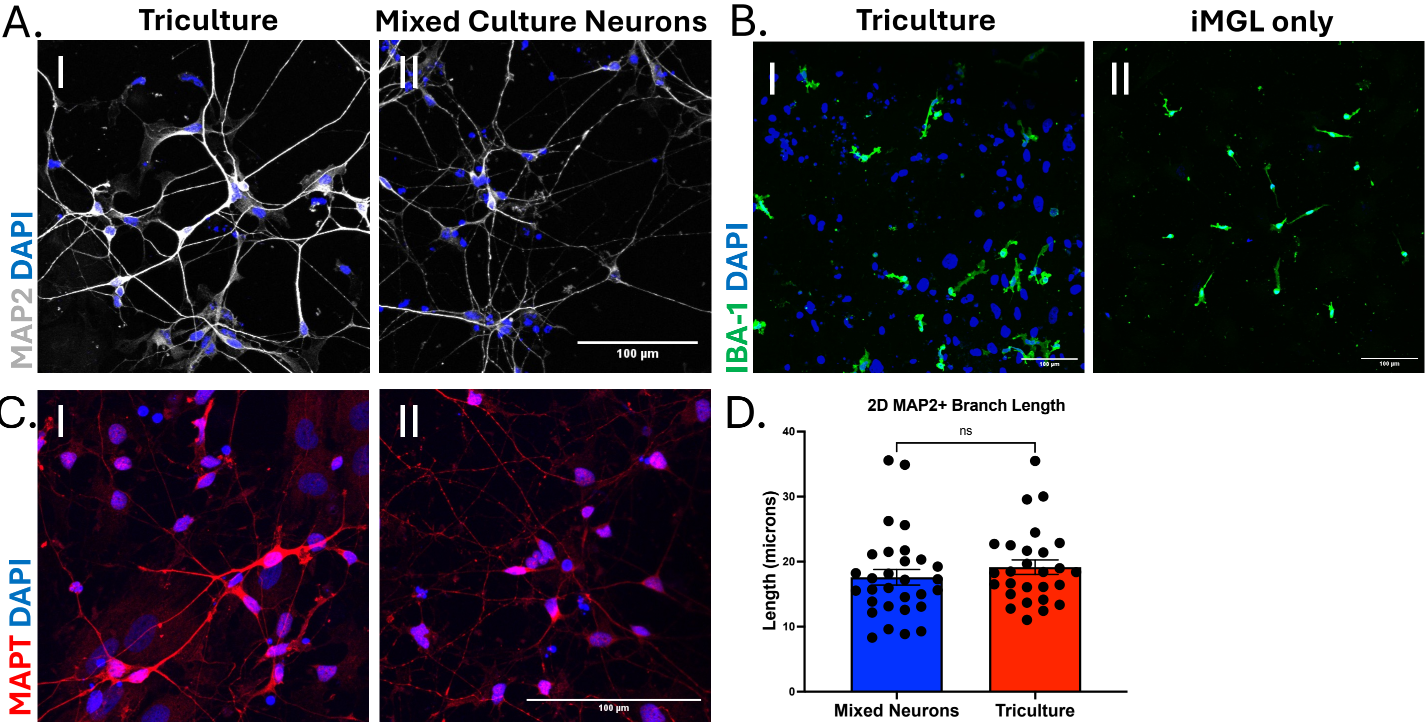


**Supplemental Figure 2. Triculture of microglia, neurons, and astrocytes results in healthier neuronal staining.** (A) Representative IF images of neuron morphology (MAP2, white) in I) tricultures containing iMGL and II) mixed culture neurons lacking iMGL. Scale bar = 100 µm. Neurons have more robust MAP2 staining in tricultures compared to mixed culture neurons. (B) Representative IF images of iMGL (IBA-1, green) in I) tricultures and II) cultures of iMGL only. Scale bar = 100 µm. iMGL have a more rod-like morphology in monocultures compared to tricultures. (C) Representative IF images of neuron morphology (MAPT, red) in I) tricultures compared to II) mixed culture neurons. Scale bar = 100 µm. Neurons have stronger MAPT staining in tricultures compared to mixed neuronal cultures. (D) Quantification of MAP2+ neuronal branch length in tricultures vs. mixed culture neurons. No significant difference in branch length was observed. Error bars, mean ± SEM. *$p < 0.05$, **$p < 0.002$, ***$p < 0.0002$, ****$p<0.0001$ by unpaired t-test.

**Supplemental figure 3.**

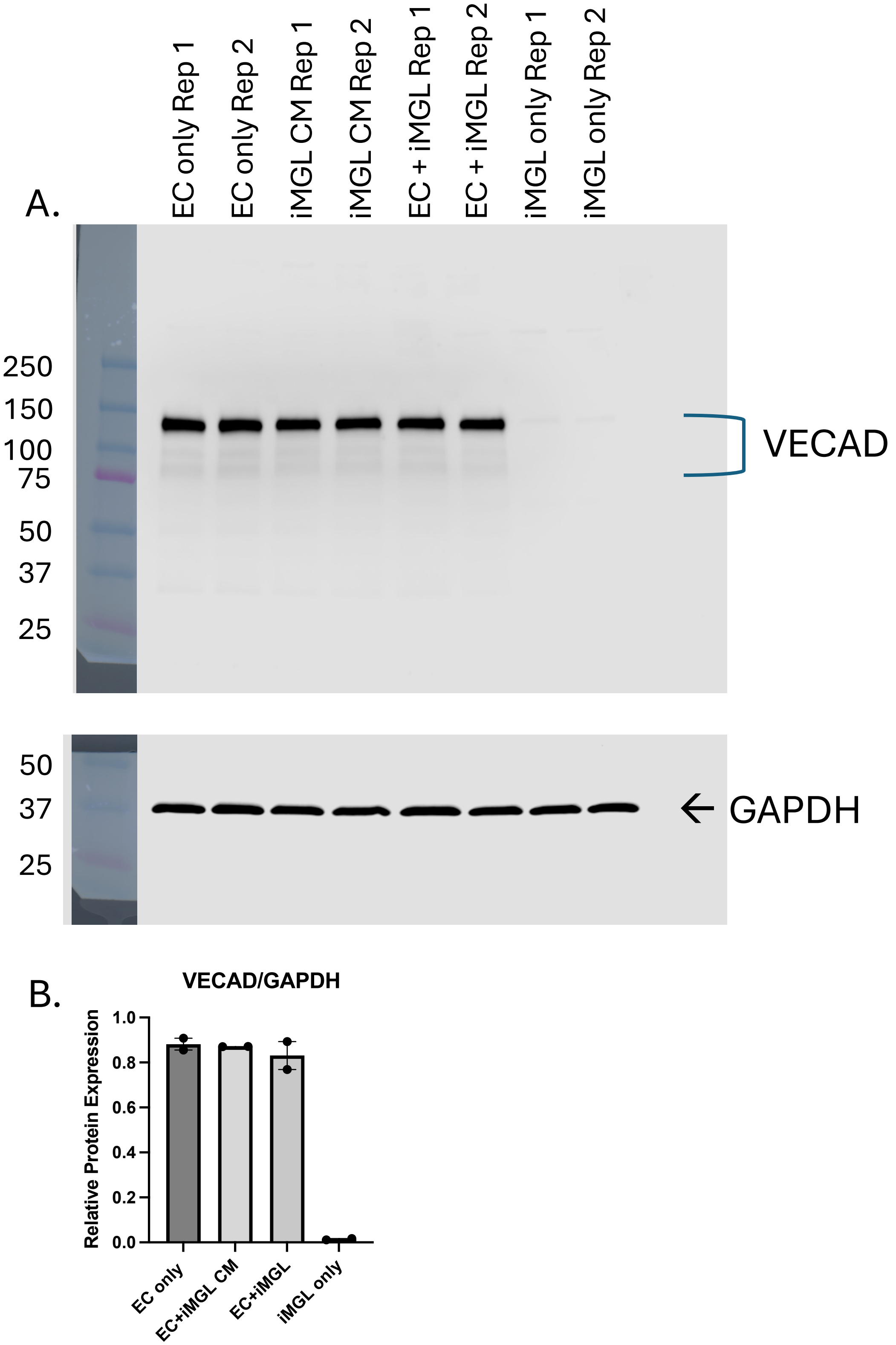


**Supplemental figure 3. Western blot for VECAD on conditioned media treated ECs and EC-iMGL co-cultures.** (A) Western blot for VE-Cadherin (VECAD) and GAPDH on protein collected from cultures of EC only, microglia conditioned media (CM) treated ECs (iMGL CM), co-cultures of ECs and iMGL (EC+iMGL) and iMGL only. VECAD levels are consistent across cultures containing ECs and there is no VECAD expression in iMGL only controls. (B) Quantification of relative VECAD expression normalized to GAPDH. Error bars, mean ± SEM.

### Supplemental Figure 4.

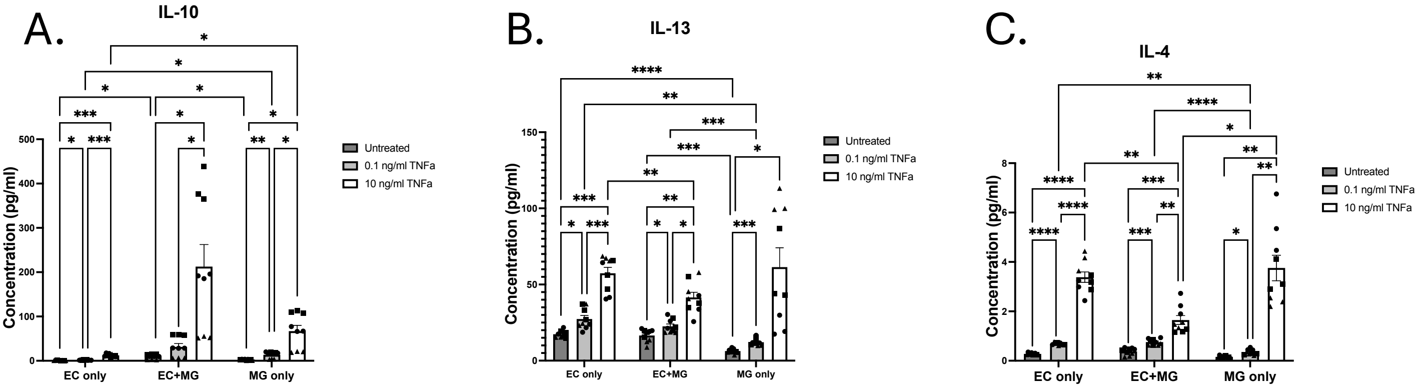


**Supplemental Figure 4. Comparison of anti-inflammatory cytokine levels between EC-iMGL co-cultures and monocultures following TNFα treatment.** (A) IL-10, (B) IL-13, and (C) IL-4 levels (pg/mL) in media from EC, iMGL, or EC-iMGL cultures, quantified by MSD immunoassay following 18 h untreated, treated with 0.1 ng/mL TNFα, or treated with 10 ng/mL TNFα. n=3 biological replicates. All Plots: mixed-effects analysis with Tukey's multiple comparisons test. Error bars, mean ± SEM. *p < 0.05, **p < 0.002, ***p < 0.0002, ****p<0.0001.

### Supplemental Figure 5.

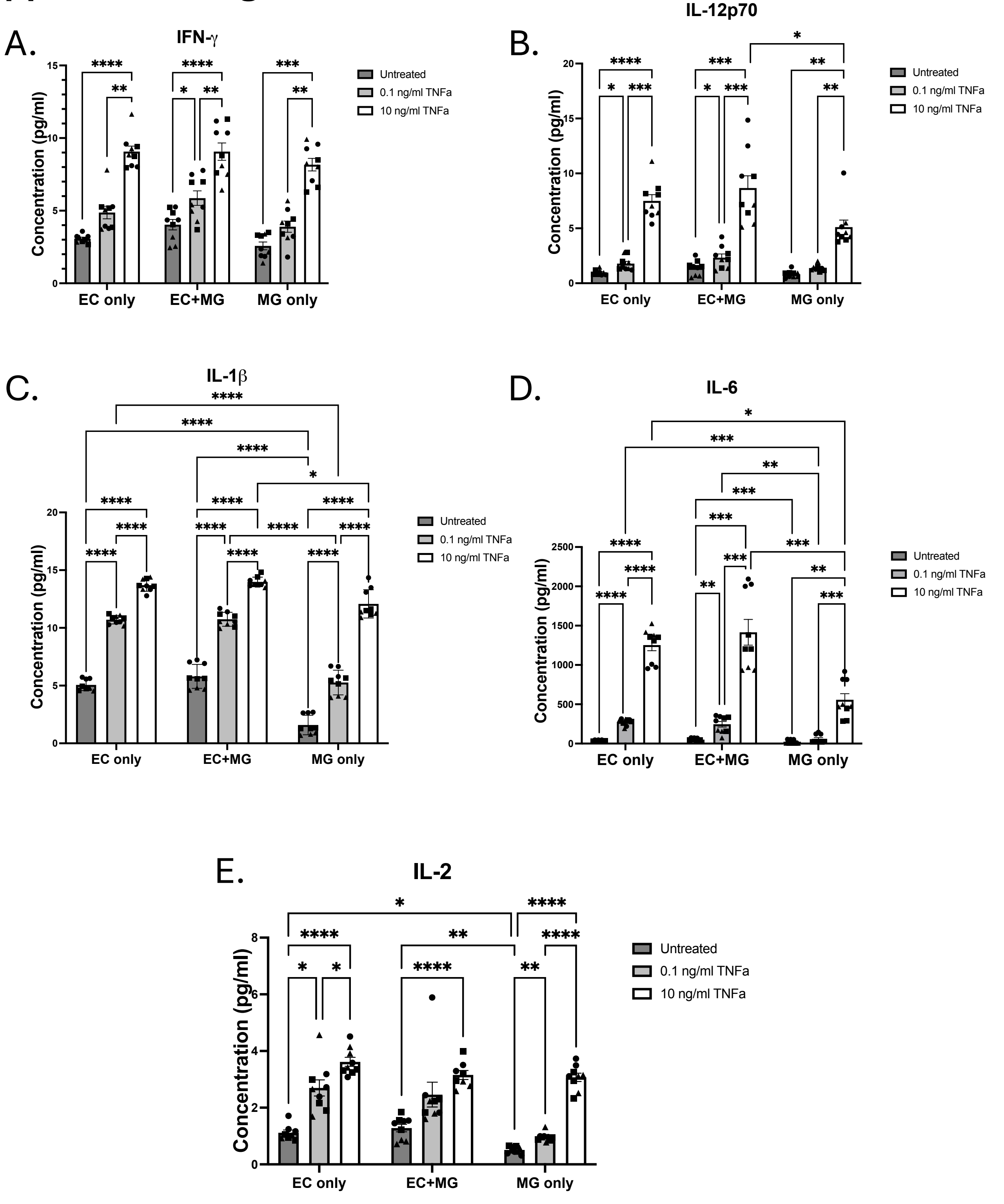


**Supplemental Figure 5. Comparison of pro-inflammatory cytokine levels between EC-iMGL co-cultures and monocultures following TNFα treatment.** (A) IFN-γ, (B) IL-12p70, and (C) IL-1β (D) IL-6 (E) IL-2 levels (pg/mL) in media from EC, iMGL, or EC-iMGL cultures, quantified by MSD immunoassay following 18 h untreated, treated with 0.1 ng/mL TNFα, or treated with 10 ng/mL TNFα. n=3 biological replicates. All Plots: Mixed-effects analysis with Tukey's multiple comparisons test. Error bars, mean ± SEM. *p < 0.05, **p < 0.002, ***p < 0.0002, ****p<0.0001.
